# gutenTAG: A MALDI-IHC processing and quality control tool

**DOI:** 10.64898/2026.11.20.767921

**Authors:** John Abbey, Pierre Bost, Leonardo Schwarz, Mengze Zhang, Christian Panse, Paolo Nanni, Natalie De Souza, Bernd Bodenmiller

## Abstract

Coupling of immunohistochemistry with MALDI mass spectroscopy (MALDI-IHC) enables targeted spatial analysis of proteins through the use of photocleavable mass tags. Upwards of 100 proteins can be imaged in a single run due to the high multiplexing power of this technology. Coupled with mass shifts in MALDI-IHC data, this scalability creates a challenge for users, as there are currently no dedicated software tools for MALDI-IHC data processing. Here we introduce gutenTAG, an open-source processing and quality control tool specifically designed for MALDI-IHC. gutenTAG uses a novel strategy based on grouping clusters of m/z bins originating from the same molecular species into single entities termed metapeaks. This strategy is robust to mass shift, generating ion images that capture the full signal of peaks corresponding to each mass tag while minimizing noise. Furthermore, we show that the metapeak strategy can be adapted to alternative open-source tools to facilitate their use in targeted imaging applications.

## Introduction

Mass spectrometry-based imaging enables spatial profiling of molecules in biological tissues. Among various mass spectrometry technologies, matrix assisted laser desorption/ionization (MALDI) has shown particular promise for a wide array of biomolecular analytes, including lipids, peptides, glycans and metabolites, achieving simultaneous detection of thousands of analytes in a single run. Direct measurement of proteins with MALDI-based imaging is currently not possible, however, due to the wide range of relevant protein sizes, low abundances of some protein targets, and challenges in identifying proteins with certainty^1^. Spatial profiling of protein in tissue is currently the domain of antibody-based methods, using cyclic fluorescence approaches such as CODEX^2^ and 4i^3^ at up to 60 plex^4,5^ and metal-based approaches such as imaging mass cytometry at 40-50 plex when applied to a single section^6^. Although powerful, these technologies are nevertheless restricted in their multiplexing capabilities due to tissue damage and autofluorescence in the fluorescence approaches and the finite repertoire of stable heavy metal isotopes available for imaging mass cytometry.

MALDI-based imaging has high mass resolving power relative to the mass detection range, is non-destructive to tissue, and has acquisition speeds fast enough to be applied to whole-slides. To harness these properties for protein imaging, MALDI-IHC was developed. Here, photocleavable mass-tags (PCMTs) that are amenable to MALDI detection are conjugated to antibodies as a reporter system^7–9^, thus enabling unprecedented levels of protein multiplexing in tissue stained with such antibody panels (**Figure 1a**). Further, in a parallel study to this one, we demonstrate that MALDI-IHC enables simultaneous, whole-slide, 133-plex protein imaging of tissue and is applicable to patient cohorts^10^.

**Figure 1.**
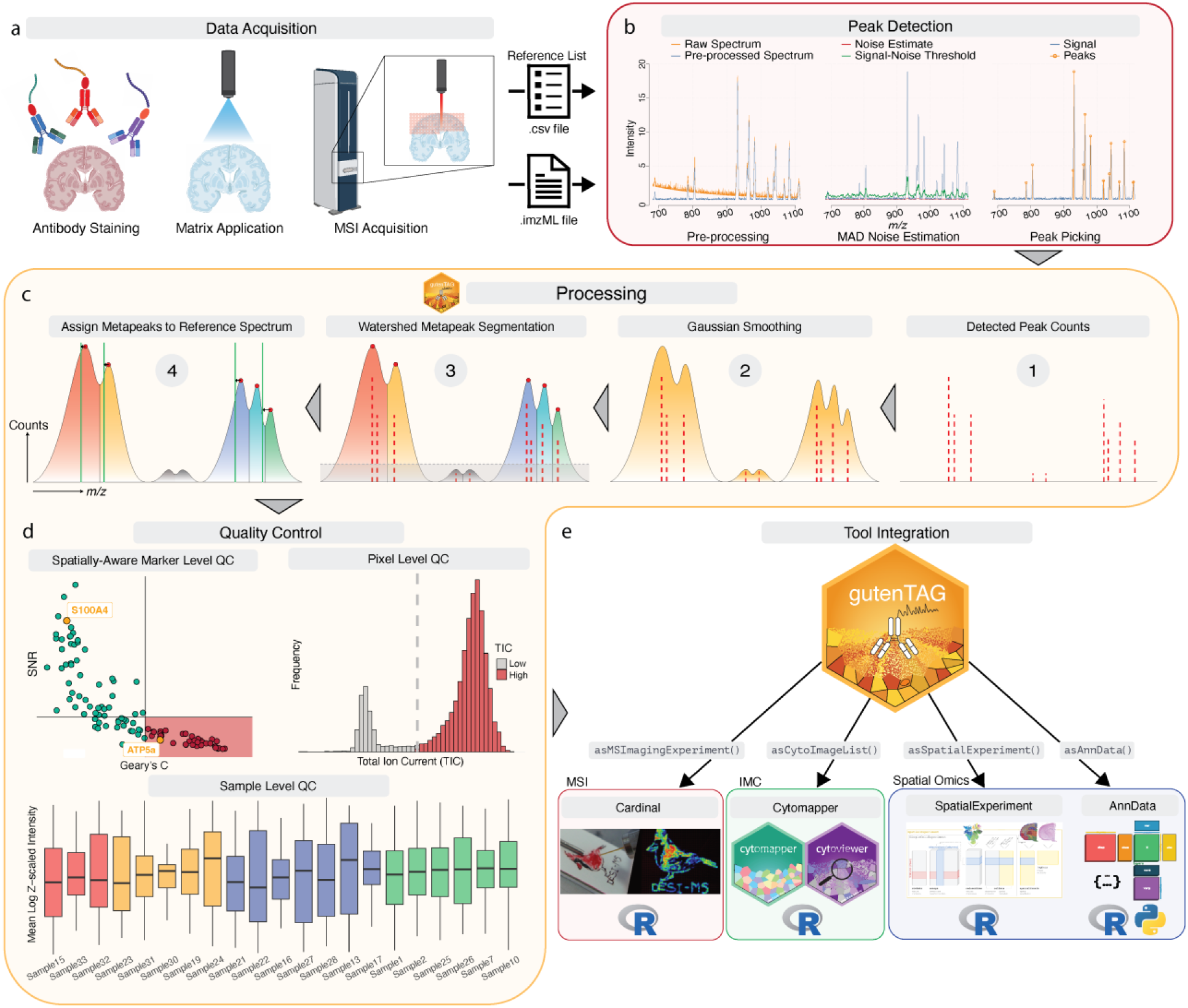
gutenTAG: An end-to-end processing and quality control pipeline for MALDI-IHC data. **a. Data acquisition process.** Tissue is stained with PCMT-conjugated antibodies, matrix is applied, and MALDI-IHC data are acquired. **b. Pre-processing and peak detection.** Raw spectral data (.imzML file) and a reference list of mass tags are loaded into R, followed by pre-processing of the raw data. A noise baseline is then estimated on the pre-processed spectra, and the SNR threshold is selected. During peak picking, any signal above this threshold is selected as a peak. **c. gutenTAG processing.** Detected peaks across all spectra are counted, creating a histogram of counts (red dashed lines) along the m/z axis. Reference list of mass tags are shown in green lines. Gaussian smoothing is then performed on the counts histogram. This is followed by watershed-based segmentation, which groups m/z bins into metapeaks. Finally, metapeaks (centers depicted by red circles) are assigned to a proximal peak in the reference list of mass tags (green lines). **d. QC.** Marker level QC filters out poor performing markers based on SNR and on spatial autocorrelation (Geary’s C). Pixel level QC removes background pixels based on TIC. Sample-level QC facilitates comparison of multiple samples/runs. **e. Tool integration.** gutenTAG’s conversion functions enable straight-forward conversion of gutenTAG data into formats compatible with various spatial data analysis frameworks.

MALDI-IHC encodes discrete molecular information (i.e., the PCMT attached to a specific target protein) into continuous mass spectra. Data-processing must then turn raw spectra at each pixel into accurately resolved molecular signals corresponding to the measured PCMTs. This is not a trivial problem as the features that enable a rich encoding of molecular information also present unique data processing challenges. First, data analysts must contend with the high dimensionality and size of the raw data. Unprocessed MALDI-IHC datasets typically contain 10^4^ to 10^5^ features (spectral bins) and 10^5^ to 10^6^ observations (pixels). The resulting raw file sizes for single whole-slide pathology samples are typically at least 10 GB, and larger file sizes will result as imaging areas are increased and pixel sizes are decreased to improve spatial resolution. This poses an infrastructural challenge, as performing even basic operations on data of this size is beyond the limits of personal computers and most virtual machines.

Second, analyzing these datasets requires distinguishing the relevant peak in a continuous mass spectrum from noise. Most features in the spectra are either background noise or peaks that do not correspond to the relevant target. One of the goals of data processing is, therefore, to discard all spectral bins except for those corresponding to the PCMTs. In principle, this could be achieved by selecting peaks only at the expected empirical m/z values of the PCMTs used in the experiment, but experimentally, the observed peaks deviate from the expected m/z location of the PCMTs due to mass shifts^9^. Thus, a third challenge in analysis of MALDI-IHC data is handling such mass shifts as well as isotopic peaks, where signal from a given peptide is distributed in regular intervals along the m/z axis.

To meet these challenges, we have developed gutenTAG, an end-to-end, open-source data processing and quality control (QC) pipeline for MALDI-IHC experiments. gutenTAG builds on the data handling capabilities of Cardinal^12,13^, an open-source R package with efficient data handling and a statistical analysis framework for mass-spectrometry-based imaging data. Cardinal is a general-purpose tool and as such does not provide dedicated algorithms for MALDI-IHC. Similarly, the open-source count-based adaptive peak calling algorithm uMAIA (unified Mass Imaging Analyzer)^14^ was designed for resolving closely neighboring peaks in lipidomics and metabolomics data, and not with the specific challenges of MALDI-IHC in mind. Distinct from other types of MSI data, MALDI-IHC data has relatively few peaks of interest, with a known but approximate (due to mass shifts) m/z location. This offered the opportunity to develop gutenTAG as a fast, lightweight tool to identify MALDI-IHC peaks with high accuracy, if mass shifts can be accounted for.

gutenTAG efficiently and accurately constructs discrete mass tag channels from continuous spectral data. At the core of its unique function is a novel metapeak generation step, where a metapeak is a single peak that groups clusters of m/z bins originating from the same molecular species. The metapeak generation step automatically accounts for mass shifts and variable peak widths and flexibly incorporates isotopic peaks via user-defined parameters, and thus effectively handling MALDI-IHC data where there are relatively few peaks of interest with a known but approximate m/z location. As an open-source tool dedicated to MALDI-IHC, gutenTAG should enable MALDI-IHC data analysis beyond users of commercial MSI analysis software.

Here, we describe the principles and operation of gutenTAG for MALDI-IHC data processing and QC, describe the effects of various parameters on metapeak generation, and detail the performance of gutenTAG relative to manually picked peaks. We show that the metapeak generation principle enhances the performance of other tools used for analysis of MALDI-IHC data. In sum, gutenTAG is an open-source data processing and QC tool designed specifically for MALDI-IHC data that enables accurate detection of peaks corresponding to PCMTs despite the presence of mass shifts and isotopic peaks.

## Results

### gutenTAG: A pipeline for MALDI-IHC data processing and quality control

Our aim in developing gutenTAG was to provide a tool specifically designed for handling the data processing challenges of antibody-based, targeted proteomic datasets such as those generated using MALDI-IHC. The gutenTAG data processing algorithm consists of two major steps: 1) data import, pre-processing, and initial peak-picking performed by Cardinal (**Figure 1a, b**) and 2) count-based metapeak generation to detect real peak centers, which are subsequently assigned to PCMTs (**Figure 1c**). gutenTAG takes as inputs a .imzML and .ibd file pair and a panel .csv file containing the information on the mass-tags and their expected m/z values (**Figure 1a**). The pre-processing step involves normalization of each spectrum, usually by total ion current (TIC), smoothing, and baseline subtraction (**Figure 1b**). This pre-processing step ensures that spectral intensities are comparable across pixels and mitigates local noise. No feature removal is performed at this stage.

Following pre-processing, we distinguish peaks corresponding to peptide mass tags from peaks that correspond to noise. Although many strategies exist for this step, we opted for a simple and stable approach in which we first estimate local noise levels and then select peaks that exceed this level. We use the median-adaptive deviation (MAD) algorithm for noise estimation; MAD calculates the median noise value within a specified m/z window, and then slides the window along the m/z axis to generate a local noise estimate. This dynamic noise estimation is beneficial as noise levels are not always constant within a spectrum. Subsequent peak picking then converts the continuous profile spectrum into discrete ‘processed’ spectra, resulting in a large reduction in the feature space. Users can choose to use Cardinal for these steps or the gutenTAG pre-processing and peak detection function, which wraps these Cardinal functions into a single pipeline.

Next, gutenTAG’s metapeak generation step follows a three-step process of 1) aggregating detected peaks into a histogram of counts, 2) segmenting metapeaks, and 3) assigning metapeaks to PCMTs in the reference spectrum (**Figure 1c**). Peak aggregation involves summing all detected peaks across all pixels to generate a histogram of counts. This histogram can be interpreted as a single spectrum representing all pixels in an image, where the x-axis is in m/z, and the y-axis is the counts of all detected peaks in each pixel. The histogram is then smoothed using a Gaussian filter that can be tuned by changing its standard deviation. This provides a smooth continuous landscape for performing metapeak segmentation. In the segmentation step, a 1-D watershed transformation is applied over the smoothed count histogram, using local minima as segmentation boundaries. This transformation groups neighboring m/z bins into a single metapeak based on proximity in m/z space. Metapeak width is determined automatically by the watershed algorithm or can be set by the user such that all metapeaks are constrained to the same width. Finally, metapeaks are assigned to PCMT peaks by comparison to a reference spectrum based on a distanced threshold, called the association threshold, that is defined by the user. The metapeak maximum, the m/z value that has the highest counts within the metapeak, is then taken as the consensus m/z location for that PCMT. Multiple metapeaks can be assigned to a single reference peak, especially if a large association threshold value is chosen, e.g. > 1 m/z. If the spatial distribution of the corresponding PCMT is known, the assignment can be made manually. Alternatively, the gutenTAG function called removeDuplicates can be used to automatically select a metapeak in the case of duplicates. If two metapeaks are assigned on one side of the peak for the reference mass tag, the closest metapeak is automatically chosen. If metapeaks are on either side of the peak for the reference mass tag, the user can select the expected direction of the mass shift, which is used in gutenTAG to select the appropriate metapeak.

Implementation of QC measures is essential for processing of MALDI-IHC data. gutenTAG performs QC at the marker, pixel, and sample levels. At the marker level, gutenTAG assesses individual ion images according to their intensities, signal-to-noise ratio (SNR), and spatial autocorrelation (Geary’s C). SNR for each marker is estimated by fitting a Gaussian mixture model to the histogram of ion counts, allowing separation of background and signal components. The ratio of the means of the two Gaussian distributions is then calculated to derive the SNR separability metric. This approach enables an automated assessment of detectability of each marker. The Geary’s C statistic is used to quantify the degree of spatial autocorrelation for each marker. Values approaching 0 indicate strong local similarity among neighboring pixels, reflecting high autocorrelation, whereas values approaching 1 indicate the absence of spatial autocorrelation. This metric therefore provides a quantitative readout of how spatially clustered or dispersed a marker pattern is within the tissue.

gutenTAG then integrates these metrics into a single summary plot that provides a visual assessment of both spatial autocorrelation and SNR for all markers in the antibody panel (**Figure 1d, Figure S1a**). The bottom right quadrant of the plot contains markers with low SNR and high Geary’s C (i.e., low spatial autocorrelation), meaning they are likely poor quality or uninformative due to poor antibody performance, low or no signal for that marker, or technical issues. Conversely, the top left quadrant contains markers with high spatial autocorrelation and high SNR that are likely to be high quality and informative. Markers in the bottom left quadrant have lower SNR than those in the top left quadrant but still have considerable spatial autocorrelation (low Geary’s C scores) and may well correspond to informative, albeit low-intensity, markers. Importantly, this plot should not be used to indiscriminately filter out markers. Even some markers within the bottom right quadrant (Survivin, for example), may show a discernable spatial pattern in the images (**Figure S1b**). Thus, this plot serves as a visual guide for users that allows identification of markers that are likely to be of high and low quality in the dataset. As a default, gutenTAG sets the SNR threshold at 3 and the Geary’s C threshold at 0.7; however, the most useful values for these metrics can vary by sample (due to differences in staining quality, tissue structure, and off-tissue components) and instrument and should thus be determined empirically.

As part of marker-level QC, the variance of each marker should also be examined, as variable markers may show informative spatial patterns. Note however that marker variance is correlated to a large extent to mean marker intensity(15, 16), (**Figure S1C**), so markers with low mean intensities can be missed if high variance is a criterion for inclusion and a strict threshold is used. This can be accounted for by regressing the standard deviation (i.e., the square root of the variance) of the markers against the mean (**Figure S1c, d**). Markers above the regression threshold are over-dispersed (i.e., they have high variance given their mean value) and are likely to be of high quality.

In pixel-level QC, gutenTAG applies a threshold to the TIC histogram, thereby discriminating between high-signal and low-signal pixels (**Figure S1e**). This QC step reveals whether there is a clear signal separation between tissue areas and background and helps identify regions of samples where no tissue is present, which typically contain low intensity pixels across all channels. In sample-level QC, gutenTAG assesses the quality of the signal from all channels for a given sample (i.e., SNR, Geary’s C, the TIC histogram, mean-variance plots, and residual plots, as described above). Any or all of these may then be compared between samples. This is of particular relevance to large cohort studies, where comparable sample intensities are imperative.

gutenTAG’s design philosophy is simple: A functional approach centered around the use of simple dataframes and lists is employed rather than an object-oriented approach. We made this choice since MALDI-IHC data are analyzed with various approaches. gutenTAG offers conversion functions that allow users to port processed data to their preferred analysis tool (**Figure 1e**). gutenTAG currently supports conversion to MSImagingExperiment (Cardinal), AnnData^17^, SpatialExperiment^18^, and CytoImageList (Cytomapper^19^/Cytoviewer^20^). The conversion function allows users to leverage a variety of tools, including image visualization, spectral analysis, spatial analysis, and multi-omic integration. In summary, gutenTAG provides a processing and QC framework that robustly creates single-marker channels for downstream spatial proteomics analysis.

**Supplementary Figure 1.**
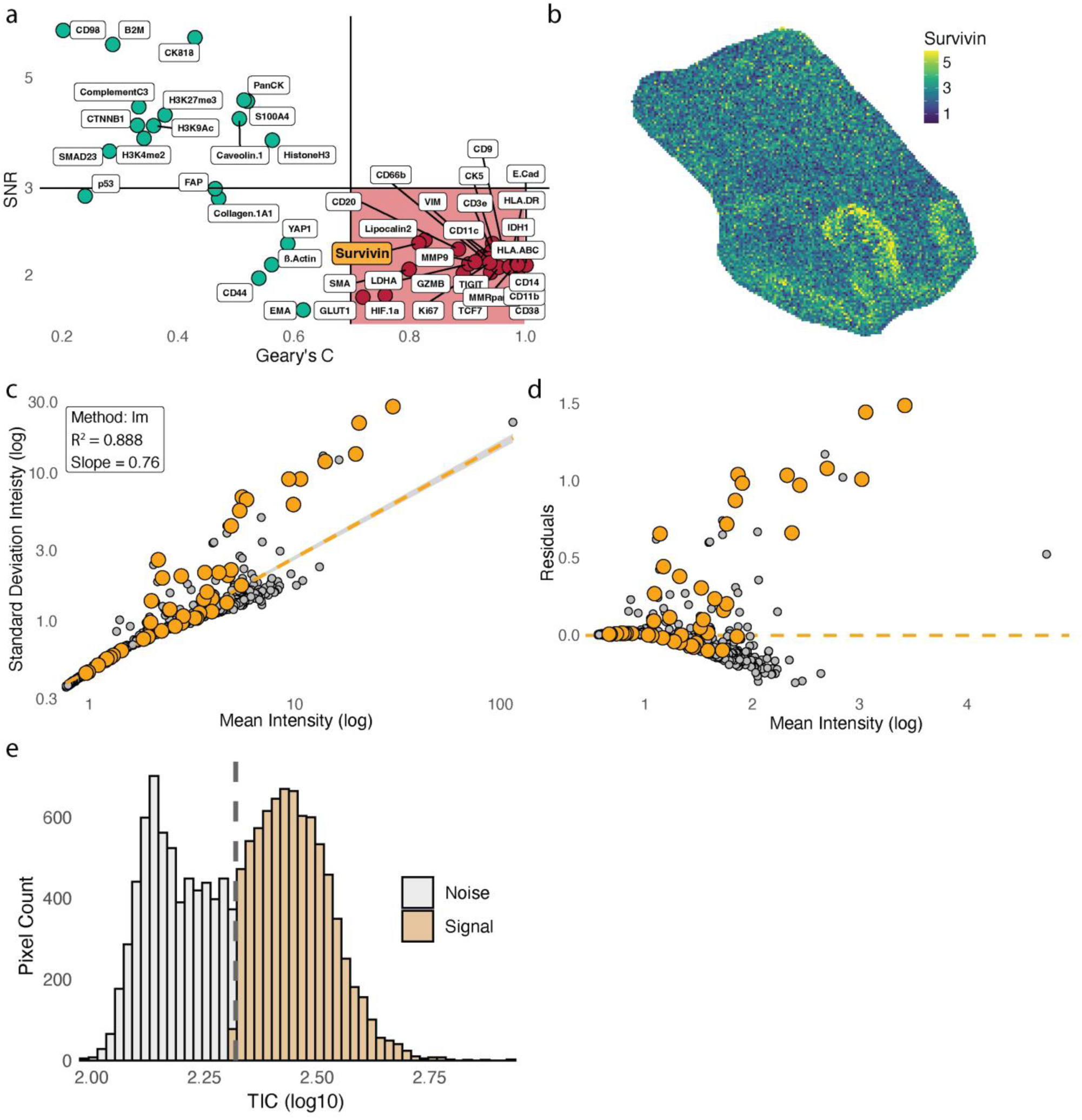
QC of data from a colon sample stained with 46-plex panel using gutenTAG. a. Scatter plot of markers according to Geary’s C spatial autocorrelation and SNR. Survivin is shaded in orange for emphasis. b. The ion image for Survivin, illustrating that even a marker within low SNR and high Geary’s C may contain some informative spatial signal. Survivin intensity is in arbitrary units. c. Scatter plot of mean metapeak intensities vs. metapeak standard deviation. Each point corresponds to the metapeak for a single marker. Orange points are assigned metapeaks, grey dots are unassigned. The orange line is the linear regression of log standard deviation against log mean. d. Scatter plot of mean metapeak intensities vs. residuals to the linear regression of mean vs. standard deviation. Each point corresponds to a single marker. The orange line is the linear regression of log standard deviation against log mean. e. Histogram of the TIC of all assigned metapeaks in the sample, separated by Otsu thresholding (grey line).

### Parameters that affect metapeak generation in gutenTAG

A number of parameters used during pre-processing and data processing control the output of gutenTAG’s metapeak generation (**Figure 2a-e**). Since the gutenTAG pipeline follows a fixed order, these parameters are interdependent, with upstream parameters influencing the effect of subsequent parameters. There are default values for all parameters, but the optimal parameter set will be instrument- and data-dependent and must be determined empirically. Here, we describe the detailed effect of each parameter on data processing using a healthy human colon tissue section stained with a 46-plex panel (**Figure S2a, b**); we further show that consistent effects are seen on other sample types, a tonsil sample stained with the same 46-panel, and an endometrial cancer sample stained with a 117-plex panel^8^ (**Figure S3**).

**Figure 2.**
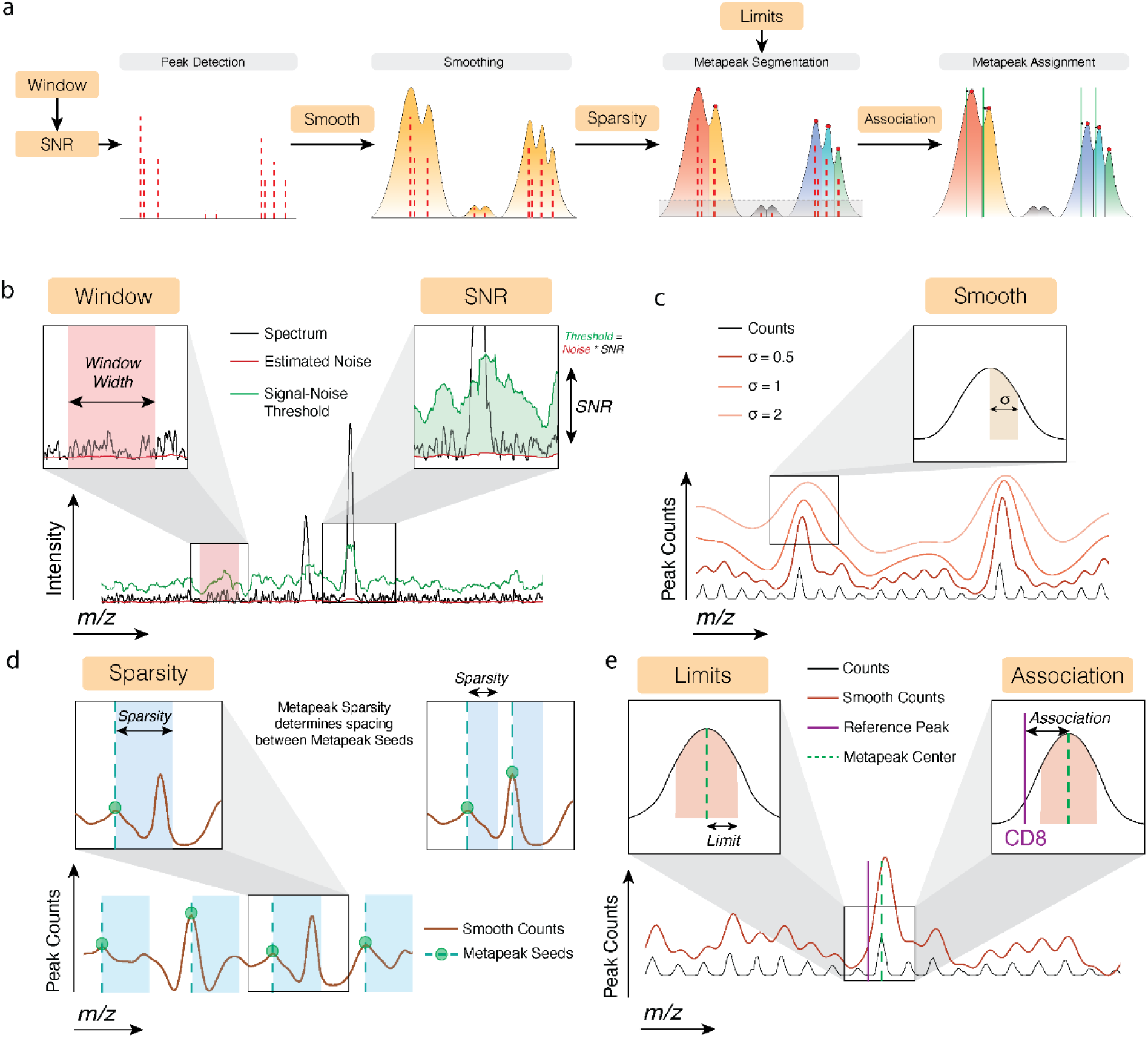
gutenTAG parameters that affect metapeak generation. a. **Parameter dependencies**. Orange boxes are labeled with tunable parameters. Arrows point toward downstream parameters and downstream metapeak generation steps that depend on upstream parameters. b. **Peak detection parameters.** The window width and SNR parameters control the output of the peak detection step. An example spectrum from a single pixel is shown. The black line shows the pre-processed spectrum, the red line is the estimated adaptive noise line, and the green line is the SNR threshold line. The window width parameter is illustrated in the left-hand zoomed window. The value of this parameter determines the width of the area from which noise is calculated as the estimator moves along the spectrum. The SNR parameter is illustrated in the right-hand zoomed window. The SNR threshold line (green line), above which signal is defined as a peak, is a function of the estimated noise line (red line) and the tunable SNR parameter (black double-sided arrow). The green shaded area represents the difference between the SNR threshold and the noise line. c. **Smoothing parameter.** Illustration of counts histogram (black line) and several smoothed counts histograms (orange lines) produced when different smoothing (σ) parameter values are used. The zoomed window in the top right indicates schematically that the degree of smoothing is controlled by σ, which determines the width of the Gaussian kernel used for smoothing. d. **Sparsity parameter.** Illustration of a smoothed counts histogram (brown line) and metapeak seeds (dashed teal lines), which are the initial locations for metapeak segmentation. Green circles correspond to local maximum values on the smoothed counts histogram. The zoomed window in the top left illustrates the use of a large sparsity parameter value, precluding placement of a second metapeak seed. The zoomed window on the top right depicts the use of a smaller sparsity parameter value, which rectifies this problem. e. **Metapeak-defining parameters.** The limits and the association parameters determine the area of signal inclusion inside a metapeak, and the maximum distance permitted for associating a metapeak with a reference mass tag, respectively. In the spectrum, the black line represents the counts histogram and the brown line depicts the smoothed counts. The dashed green line shows the metapeak center and the solid purple line shows the reference mass tag m/z value. The limits parameter (top left) is the m/z distance on either side of the metapeak center from which signal is included in the metapeak. The association parameter (top right) determines how near a metapeak center the reference mass tag (in this case for CD8) must be in order to be assigned to it.

The *window width parameter (window)* is the width of the m/z window used for noise estimation during peak detection (**Figure 2b)**. The value for this parameter has a considerable impact on the counts histogram. Values close to 1 correspond to a narrow window for noise estimation, resulting in detection of many more peaks than would be observed with a higher value. The counts histogram generated with a window close to 1 will likely include spurious peaks that are further from the m/z of the corresponding reference mass tag (**Figure 3a, b**). Higher values correspond to a wider window (in this case 1-50 m/z was tested) for noise estimation and result in fewer detected peaks and better discrimination of true peaks from noise. For processing of the colon tissue image data, a window value of 50m/z was used, as we wanted a counts histogram with little noise for this analysis.

**Figure 3.**
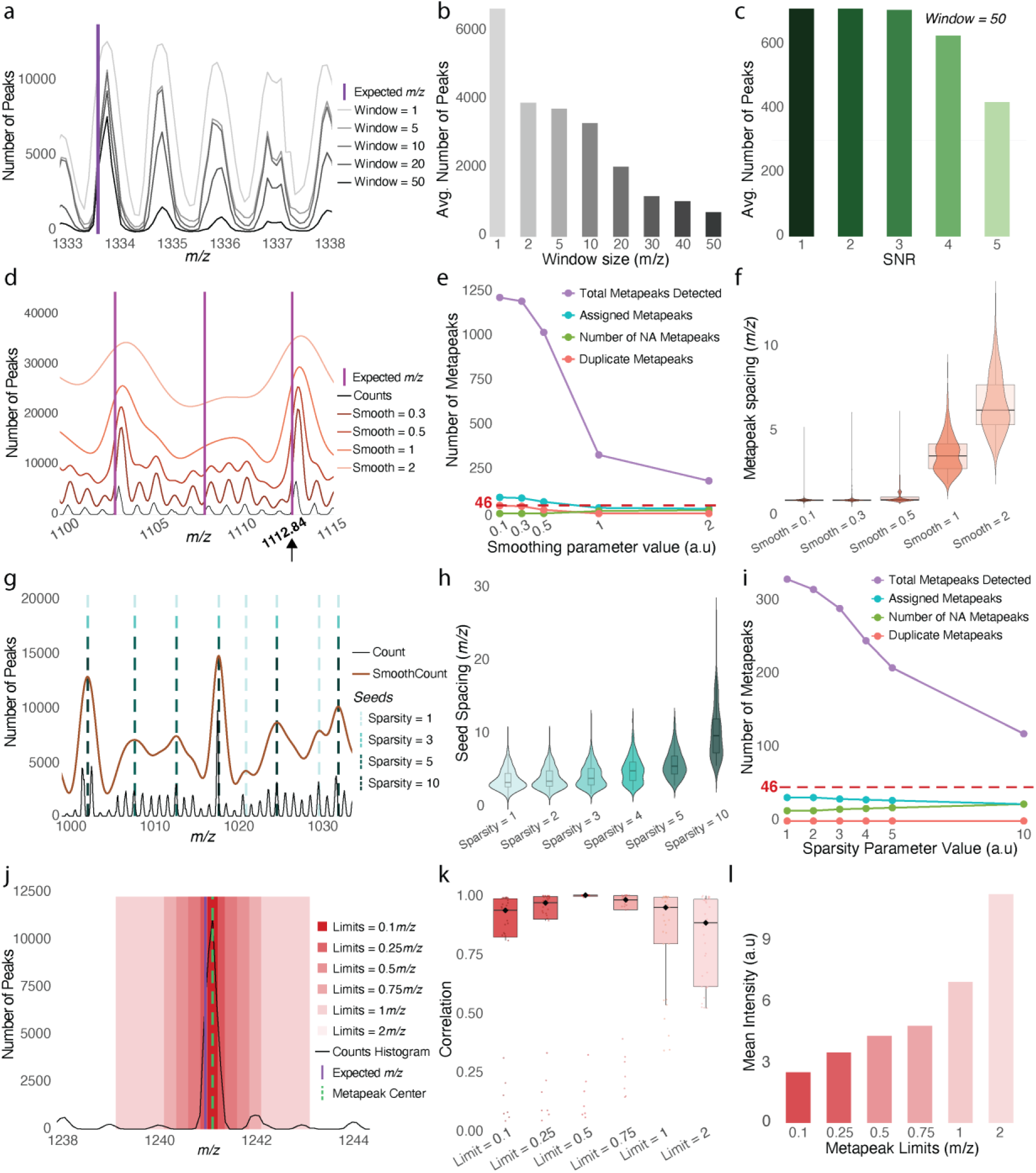
The effect of metapeak parameters on peak picking for a colon dataset. All panels show data corresponding to the image in Figure S2a. a. The effect of the window parameter on the counts histogram of detected peaks. Black and grey lines depict counts histograms corresponding to the indicated window parameter values used during peak detection. The purple line indicates the expected m/z location of a reference mass tag from the marker panel. b. Average number of detected peaks (i.e., counts) for the indicated window parameters values. c. Average number of detected peaks (i.e., counts) for the indicated SNR parameter values. d. The effect of the smoothing parameter on the smoothed counts landscape. The black line indicates the counts histogram. Brown and orange lines depict smoothed count histograms for different smoothing parameter values. The purple lines indicate the expected m/z locations of reference mass tags from the marker panel. e. Number of metapeaks detected with different smoothing parameter values. Plotted are the total number of metapeaks (purple), the number of assigned metapeaks (cyan), the number of duplicate metapeaks (red), and the number of NA metapeaks (no metapeak for a given PCMT, green) detected for the indicated smoothing parameter values. f. Average spacing of metapeaks on the m/z axis for different smoothing parameter values (a.u.). g. The effect of the sparsity parameter on the location of metapeak seeds. The black line indicates the counts histogram of detected peaks, and the brown line indicates the smoothed counts histogram. Dashed lines indicate the locations of metapeak seeds. Light shades of cyan indicate seeds from low sparsity parameter values, and darker shades indicate seeds from higher values. h. Average spacing of seeds on the m/z axis for different sparsity parameters (a.u.). i. Number of metapeaks detected with different sparsity parameter values. Plotted are the total number of metapeaks (purple), the number of assigned metapeaks (cyan), the number of duplicate metapeaks (red), and the number of NA Metapeaks (green) detected for the indicated sparsity parameter values. j. The effect of the limits parameter on the signal integrated into metapeaks. The purple line indicates the expected m/z location of a reference mass tag from the marker panel. The dashed green line marks the center of a given metapeak. The black line is the counts histogram of detected peaks. Red rectangles are metapeak limits for varying parameter values. Darker shades of red indicate signal integration areas for narrower limits, and lighter shades indicate the area for wider limits. k. Average correlation of single channel images to a manually picked image for a given marker for a series of metapeak limit values. l. Mean intensity of metapeaks for a series of metapeak limit values.

The *SNR threshold* is the SNR level above which a spectral signal is considered to be a peak. During peak detection, this threshold is determined by multiplying the estimated noise level by the SNR (**Figure 2b**). Increasing the SNR threshold results in fewer detected peaks (**Figure 3c, Figure S2c**). The number of detected peaks is influenced by both the SNR threshold and the chosen window value. For analysis of the colon dataset, we chose an SNR of 2, as we wanted to detect some PCMTs expected to have low signal. In general, the SNR threshold should be chosen to minimize the inclusion of noise while retaining low intensity signals as much as possible.

The *smoothing parameter* is the standard deviation of the Gaussian kernel used for smoothing the counts histogram. Smoothing is performed before watershed segmentation to suppress spurious local minima and provide a smoother landscape for segmentation. Low values of the smoothing parameter result in a smoothed histogram that does not differ much from the unsmoothed counts histogram, whereas higher values smooth over local structures (**Figure 2c**). In the colon image analysis, a smoothing value of 0.3 resolved the isotopic envelope emanating from the monoisotopic peak at 1112.84 m/z, whereas values of 1 and 2 smoothed this structure into a single large peak (**Figure 3d**). Importantly, the smoothing parameter affects how many metapeaks are detected downstream. Increasing the smoothing parameter results in fewer overall metapeaks, fewer assigned metapeaks (**Figure 3e**), and greater metapeak-to-metapeak spacing (**Figure 3f**). Decreasing the smoothing parameter results in more duplicate metapeaks, which requires downstream removal of duplicates, and fewer unassigned reference peaks (i.e., NA Metapeaks). The ideal smoothing value is one that minimizes the number of both duplicate metapeaks and NA Metapeaks. In the colon dataset, a smoothing parameter of 0.3 is a good choice if the user wants metapeaks for each of the individual peaks in the isotopic envelope at 1112.84 m/z; however, a value of 1 or 2 would be better if the user wants to incorporate the signal from all isotopic peaks into a single metapeak.

The *sparsity parameter* controls the spacing of seeds during watershed segmentation of metapeaks. Local maxima on the smoothed counts histogram determine the location of these seeds (**Figure 2d**). Increasing sparsity values result in larger spacing between seeds and fewer total metapeaks (**Figure 3g, h**). This is reflected in the number of total metapeaks and the number of assigned metapeaks, with low sparsity values resulting in more metapeaks than higher values (**Figure 3i**). Importantly, where there is a seed from a high sparsity value, there is always a lower sparsity value at the same location (**Figure 3g**). The choice of sparsity value will depend on whether users wish to resolve neighboring peaks, such as individual isotopic peaks (in this case, a small value, e.g., 1, should be chosen) or only the most prominent monoisotopic peak (e.g., a sparsity parameter of 3 should be chosen). The choice of the sparsity parameter should be informed by knowledge about mass tag spacing. For instance, if you expect mass tags to be spaced 5 Da apart, the sparsity parameter should be less than 5.

The *limits* parameter determines the m/z distance on either side of the metapeak center and thus the signal that is integrated into the metapeak (**Figure 2e**). The limits parameter can either be set by the user for each sample or determined automatically per-peak by the watershed segmentation algorithm. The latter allows for dynamic, flexible, and adaptive incorporation of signal into metapeaks but is unstable in high noise regions, resulting in excessively wide metapeaks. Furthermore, when comparing samples between groups, for instance in a study of patient cohorts, automatic selection of the limit parameter can introduce unwanted batch effects. This can be mitigated by setting manual metapeak limits. Low limit values (e.g., 0.1 m/z) can result in only a partial coverage of the peak, whereas high values can lead to inclusion of signal from neighboring peaks (**Figure 3j**). The latter is undesirable, as this may capture an incorrect spatial distribution (**Figure 3k**). Increasing the limits parameter also increases the average signal intensity for metapeaks (**Figure 3l**). For the colon dataset, a limits value of 0.5 m/z maximizes peak coverage but does not include noise from neighboring peaks.

The *association parameter* controls the maximum permitted m/z distance for associating a metapeak with a particular PCMT (**Figure 2e**). Metapeaks within this distance are assigned to the corresponding PCMT, and those outside are considered unassigned (**Figure S2e)**. High values for this parameter result in more assigned peaks as well as duplicate peaks (**Figure S2d**). Achieving higher levels of peak assignment by increasing the association parameter is inadvisable as many assigned peaks are likely to be false positives. We recommend choosing association parameter values based on the mass shift in the data. Higher association values are suitable only when mass shifts are substantial. Since mass shifts are highly instrument-dependent, it is essential that this parameter be determined empirically. A conservative choice for the association parameter greatly mitigates false positive peak assignment. For the colon dataset, we used an association parameter of 0.5 m/z either side of the reference mass tag, as the most extreme mass shift we observed in the dataset was just below 0.5 m/z away from the target. The relatively sparse MALDI-IHC spectrum and the fact that the approximate m/z locations of specific peaks are known makes this feasible in this context, whereas in untargeted settings such values would be less appropriate.

**Supplementary Figure 2.**
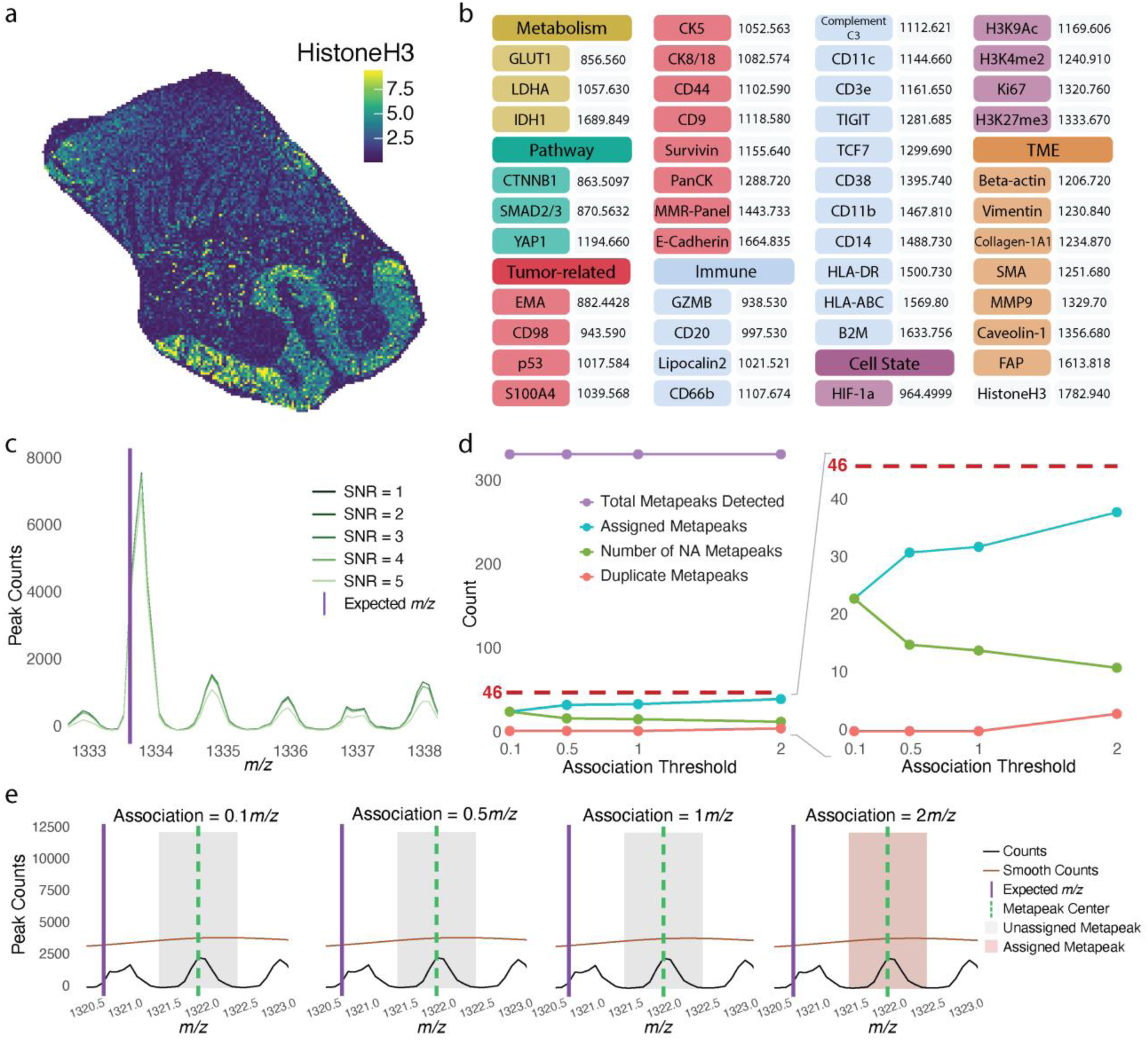
The effects of metapeak parameters on peak picking. a. Single channel image for Histone-H3 mass tag. b. List of targets of the 46-plex panel used to stain colon sample (left) and expected m/z value of the mass tag (right). c. The effect of the SNR parameter on the counts histogram of detected peaks. Peak counts vs. m/z value with increasingly bright lines indicative of mean spectra from increasing SNR threshold values. d. Number of metapeaks detected with different association threshold values. Plotted are the total number of metapeaks (purple), the number of assigned metapeaks (cyan), the number of duplicate metapeaks (red), and the number of NA metapeaks (green) for the indicated association parameter values. Lower count regions are magnified on the right. e. Example of aberrant metapeak assignment with large association value. The black lines indicate counts histograms. The brown lines depict smoothed count histograms. The purple lines indicate the expected m/z location of a reference mass tag. Unassigned and assigned peaks are shown at the indicated association parameter values. As seen on the far right plot, a metapeak far from the reference mass tag can be erroneously assigned to that tag using an association parameter value of 2 m/z.

**Supplementary Figure 3.**
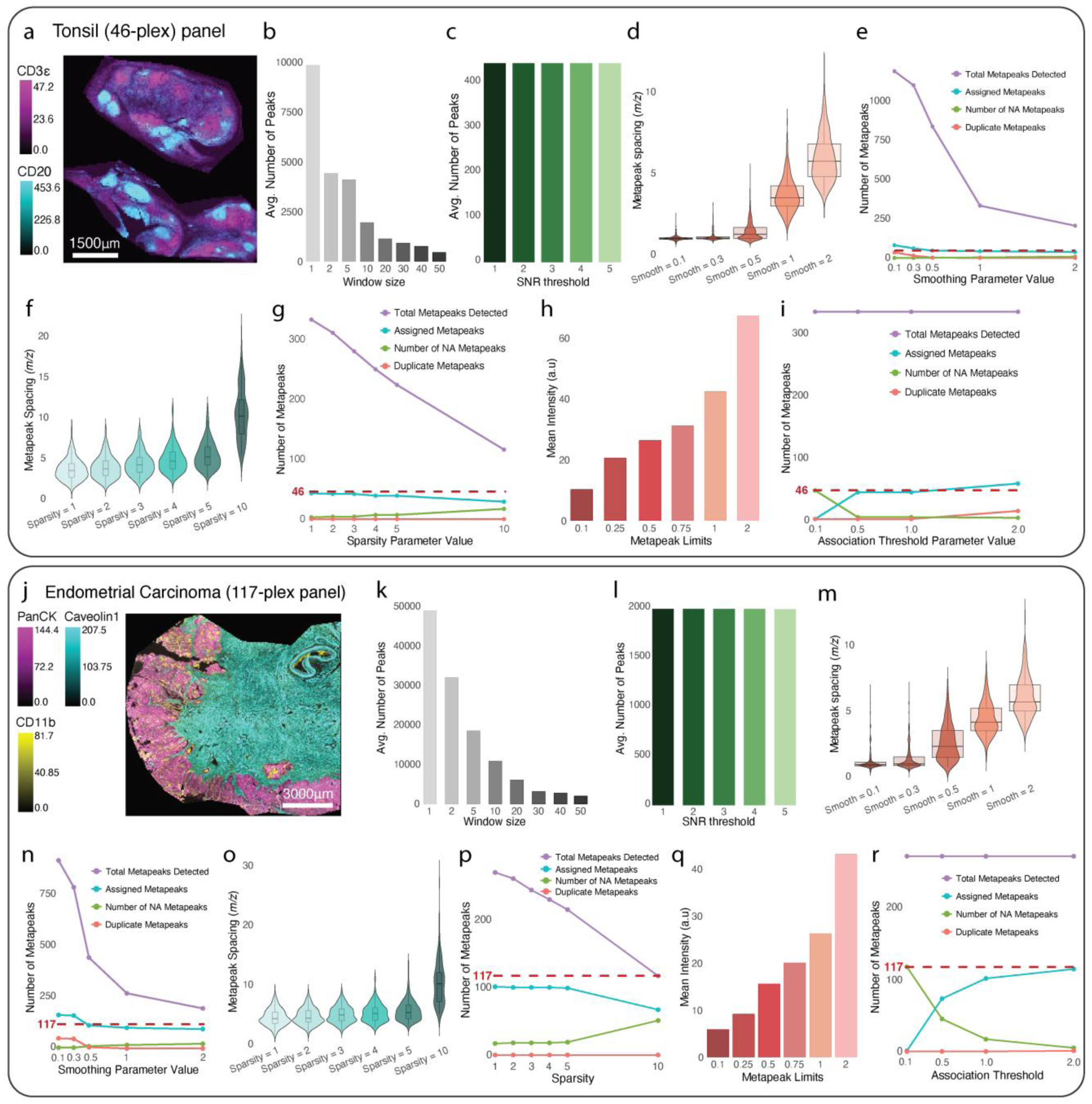
Metapeak parameter effects on data processing are consistent across multiple samples. a. Tonsil sample stained with 46-plex panel. b. Average number of detected peaks (i.e., counts) for the indicated window parameters values. c. Average number of detected peaks (i.e., counts) for the indicated SNR parameter values. d. Average spacing of metapeaks on the m/z axis for different smoothing parameter values (a.u.). e. Number of metapeaks detected with different smoothing parameter values. Plotted are the total number of metapeaks (purple), the number of assigned metapeaks (cyan), the number of duplicate metapeaks (red), and the number of NA metapeaks (no metapeak for a given PCMT, green) detected for the indicated smoothing parameter values (a.u.). f. Average spacing of seeds on the m/z axis for different sparsity parameters (a.u.). g. Number of metapeaks detected with different sparsity parameter values. Plotted are the total number of metapeaks (purple), the number of assigned metapeaks (cyan), the number of duplicate metapeaks (red), and the number of NA Metapeaks (green) detected for the indicated sparsity parameter values (a.u.). h. Mean intensity of metapeaks for a series of metapeak limit values. i. Number of metapeaks detected with different association threshold values. Plotted are the total number of metapeaks (purple), the number of assigned metapeaks (cyan), the number of duplicate metapeaks (red), and the number of NA metapeaks (green) for the indicated association parameter values. Lower count regions are magnified on the right. j. Endometrial cancer sample stained with a 117-plex panel. k. Average number of detected peaks (i.e., counts) for the indicated window parameters values. l. Average number of detected peaks (i.e., counts) for the indicated SNR parameter values. m. Average spacing of metapeaks on the m/z axis for different smoothing parameter values. n. Number of metapeaks detected with different smoothing parameter values. Plotted are the total number of metapeaks (purple), the number of assigned metapeaks (cyan), the number of duplicate metapeaks (red), and the number of NA metapeaks (no metapeak for a given PCMT, green) detected for the indicated smoothing parameter values (a.u.). o. Average spacing of seeds on the m/z axis for different sparsity parameters (a.u.). p. Number of metapeaks detected with different sparsity parameter values. Plotted are the total number of metapeaks (purple), the number of assigned metapeaks (cyan), the number of duplicate metapeaks (red), and the number of NA Metapeaks (green) detected for the indicated sparsity parameter values (a.u.). q. Mean intensity of metapeaks for a series of metapeak limit values. r. Number of metapeaks detected with different association threshold values. Plotted are the total number of metapeaks (purple), the number of assigned metapeak (cyan), the number of duplicate metapeaks (red), and the number of NA metapeaks (green) for the indicated association parameter values. Lower count regions are magnified on the right.

### Comparison to alternative methods

Next, we evaluated gutenTAG’s performance compared to that of alternative open-source methods, testing for speed and accuracy; as a “ground truth” reference we used manually picked peaks. We compared gutenTAG to uMAIA, a Python-based tool developed for processing lipidomics imaging data, and to a simple binning method in which m/z values within increasing limits (0.1, 0.2, 0.5 and 1 m/z) were grouped into bins corresponding to peaks from mass tags in the reference panel. This analysis was performed on the colon tissue sample stained with a 46-plex marker panel (**Figure S2b**).

To evaluate the run-time of these three approaches, we generated a series of datasets from the colon image analysis containing between 10 and 100,000 pixels. In each case, raw data were pre-processed identically (TIC normalization, spectral smoothing, and baseline reduction), and the number of marker channels were also identical. The gutenTAG pipeline included peak detection, metapeak segmentation, and assignment of metapeaks to the mass tags in the reference panel. The simple binning approaches were performed using Cardinal and consisted of four steps. First, profile spectra were recalibrated so that peaks across pixels were aligned, followed by peak picking using MAD-based noise estimation with the same parameters as in gutenTAG. The picked spectra were then aligned to the reference peaks, and, finally, the aligned spectra were binned with a range of binning widths (0.1, 0.2, 0.5 and 1.0 m/z). For uMAIA analysis, we ran the uMAIA.pf.run() function for each of the pre-processed datasets. The parameters were chosen to align as closely as possible with the gutenTAG parameters. For all datasets, the processing pipelines were run three times on a single core of an AMD EPYC 9554 processor without any parallelization. The average runtime and the standard error were recorded in each case. (**Figure 4a**). As expected for this relatively simple process, the binning approach was faster than uMAIA, and the run-time of gutenTAG was comparable to simple binning.

**Figure 4.**
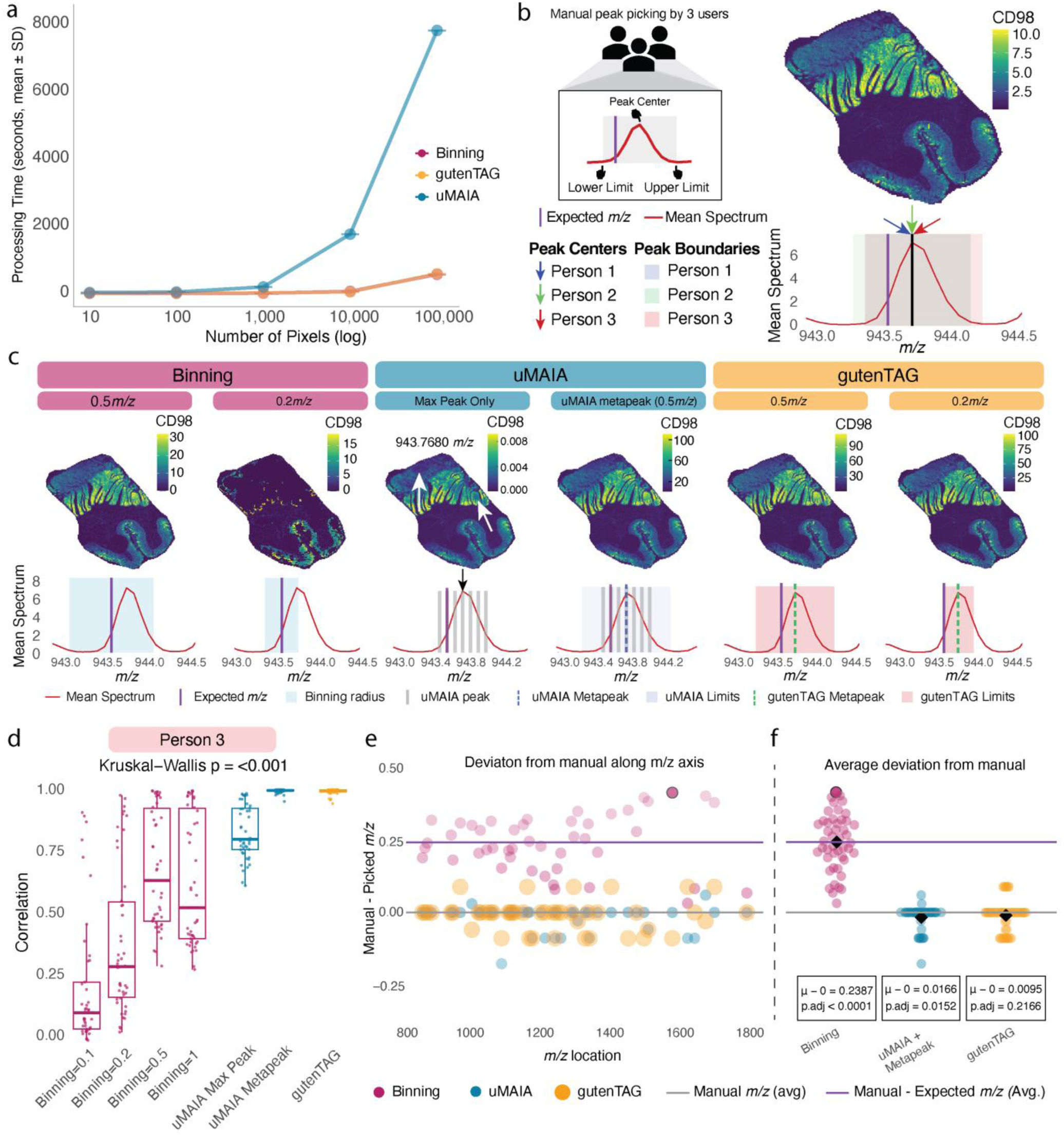
Performance comparison of gutenTAG to alternative methods. Multi-user manually picked peaks (**Figure S3**) were used as the ground truth. a. Run times of the indicated methods on datasets of increasing size (i.e., pixel number). Comparisons were conducted using a single core. b. Generation of ground truth for the methods comparison. Top left: Manual peak picking was performed independently by three users. Each individual selected a peak center and lower limit and upper limit to best represent the boundaries of that peak. This was done for all peaks corresponding to mass tags in the reference panel. Top right: A manually picked example image at 943.590 m/z, the m/z for the CD98 PCMT. Bottom: The mean spectrum of the peak picked for CD98 by the three users (red) and the expected m/z location of the CD98 tag (purple line). Blue, green, and red arrows point to the peak centers picked by individual users, and the corresponding rectangles are manually picked boundaries. c. Ion images for CD98 with reference m/z of 943.590 (upper) and corresponding mean spectra across all pixels (lower) for simple binning (magenta) with limits of 0.5 m/z and 0.2 m/z, for UMAIA (blue) taking only the peak with the highest intensity (Max Peak Only) or for the aggregation of all peaks within 0.5 m/z limits (uMAIA metapeak), and for gutenTAG (yellow) with limits of 0.5 m/z and 0.2 m/z around the metapeak center. The red dashed boxes indicate a lower intensity region not captured well by the uMAIA Max peak approach. d. Pixel-wise correlations between signal intensities in images for each peak picking method and signal intensities in manually picked images for the same channel. Correlations were computed for all channels. Shown here are correlations with intensities in manually picked images from a single individual (person 3). See **Figure S5d** for correlations to manually picked images from persons 1 and 2. e. Difference between manually picked m/z locations and those of peaks picked with the indicated methods. For simple binning, 0.5 m/z limits were used. For the other two methods, parameters were chosen to maximize correlations to the manual images (**Figure 4c, Figure S5c**). The most extreme mass shift is highlighted with a black border. f. Average difference between manually picked m/z locations and those of peaks picked with the indicated methods. The *p* values were generated using Benjamini-Hochberg-corrected *t*-tests. The most extreme mass shift is highlighted with a black border.

Next, to evaluate peak picking accuracy, we compared the outputs of each method to manual peak picking. To generate this manual ground truth, three individuals selected peaks at m/z locations at or near the expected m/z location for a PCMT, identifying both the maximum intensity value of the peak and its boundaries on either side (**Figure 4b; Figure S6a, b**). We qualitatively determined the imaging output of each method by comparison to the ground truth data. We observed, in the case of CD98, that a binning width of 0.5 m/z captured the expected ion image, whereas a width of 0.2 m/z did not as it was insufficient to cover signal from the entire peak (**Figure 4c**, left). This was expected, as simple binning takes the expected m/z values of a given PCMT as the peak center and is therefore sensitive to mass shifts. gutenTAG did capture the expected ion image with binning widths (determined by the limit parameter) of both 0.5 m/z and 0.2 m/z (**Figure 4c**, right).

For uMAIA, we tested a range of smoothing values (0.1 - 1 m/z) and found that low values (0.1, 0.2, and 0.5) over-segmented the uMAIA-picked peaks, resulting in many partially complete ion images (**Figure S5a-d**). This was likely due to differing tissue heights on the slide. Conversely, at high smoothing values some peaks were not identified at all, as was the case for the mass tag for CD98 (**Figure S5e, f**). For both the smoothing and the count threshold parameters, we were unable to find a parameter set for uMAIA that detected a single peak for each PCMT in the dataset without missing expected peaks. Therefore, we chose to evaluate uMAIA parameter sets that tended to over segment peaks. We used smoothing values of 0.1 to 0.2, as these values detected the most peaks and had the highest correlation to manual picking (**Figure S5g, S6c**), and a count threshold value of 10000, as lower tested values were not sufficiently sensitive to changing the smoothing value (**Figure S5h)**. Images corresponding to uMAIA peaks with the highest local intensity typically captured the salient features of manually picked images (**Figure 4c**). Overall, signals were weak (which could pose a problem for markers with lower SNR than CD98), and areas of low intensity were sometimes missed. These differences were likely due to height differences in the tissue on the slide during acquisition, which cause a slight shift of some of the signal along the m/z axis; these signals can be missed when focusing only on the highest intensity peaks. We therefore assessed whether applying a metapeak-like approach in uMAIA similar to that used in gutenTAG could achieve higher fidelity peak picking. We selected the uMAIA peaks with the highest intensities as the central peaks and integrated neighboring peaks within signal inclusion limits (0.5 m/z). The resulting image captured the expected signal distribution (**Figure 4c**), and intensities were comparable to those of the manually picked image (**Figure 4b**).

Next, we quantitatively evaluated the output of each method relative to that of manual peak picking. First, we computed the pixel-wise correlations for all channels in the images generated with each approach to those generated from peaks manually picked by three individuals. We found that the binning approach yielded significantly lower correlations to manually picked images for all binning widths compared to gutenTAG and uMAIA (**Figure 4d, Figure S6d**). Interestingly, both gutenTAG and uMAIA resulted in very good pixel-wise correlations to manually picked images when using the metapeak approach. Using the maximum intensity uMAIA peaks yielded on average lower correlations to manually picked images than metapeak-based approaches, although the performance was better than simple binning.

Finally, we assessed how well the peaks picked by our three methods lined up with manually picked peaks along the m/z space axis. To do this, we computed the m/z distances between the manually picked peak centers and the peak centers determined by each method for all channels (**Figure 4e, f**). Peaks identified by simple binning significantly deviated from manually-picked peaks as expected, since simple binning takes the expected m/z value as the peak center and is therefore not robust to mass shifts. In contrast, peaks identified by gutenTAG and uMAIA aligned well with the manually picked peaks (**Figure 4f**), indicating that these metapeak-based approaches are robust to mass shifts in the data. Taken together, our data show that gutenTAG accurately detects peaks corresponding to PCMTs, is robust to mass shifts in the data, and has comparable runtime to simpler, less performant methods.

**Supplementary Figure 4.**
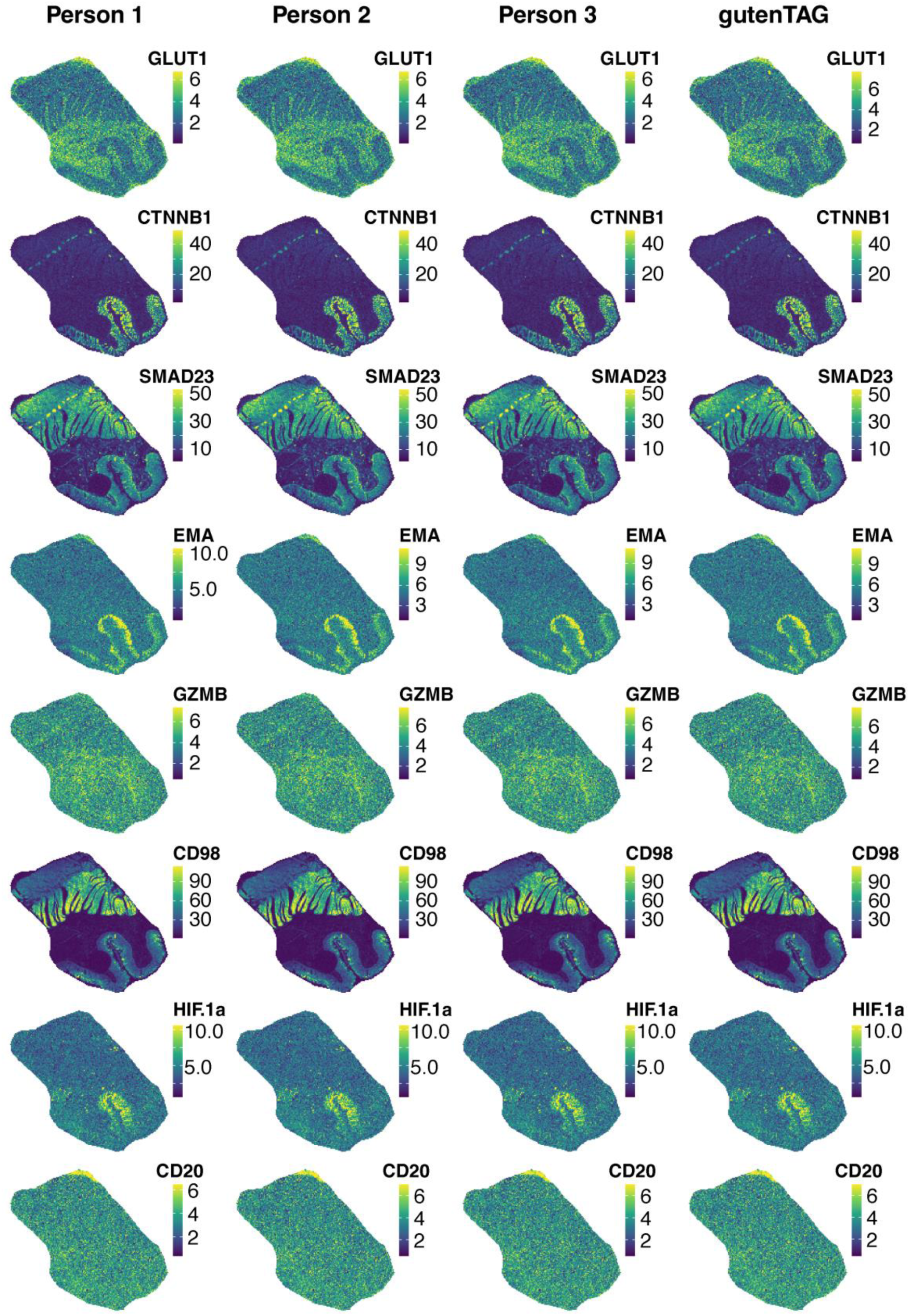
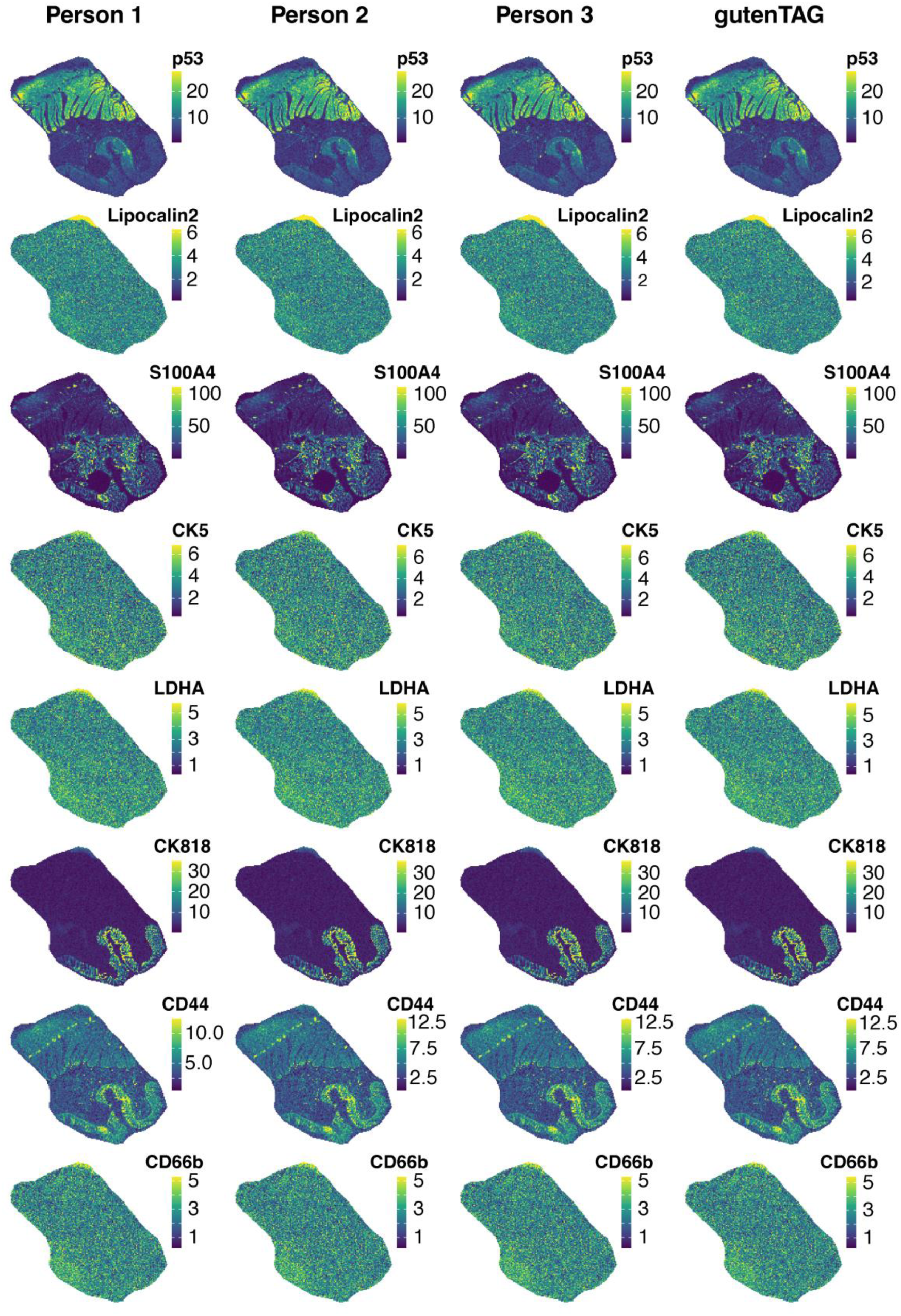
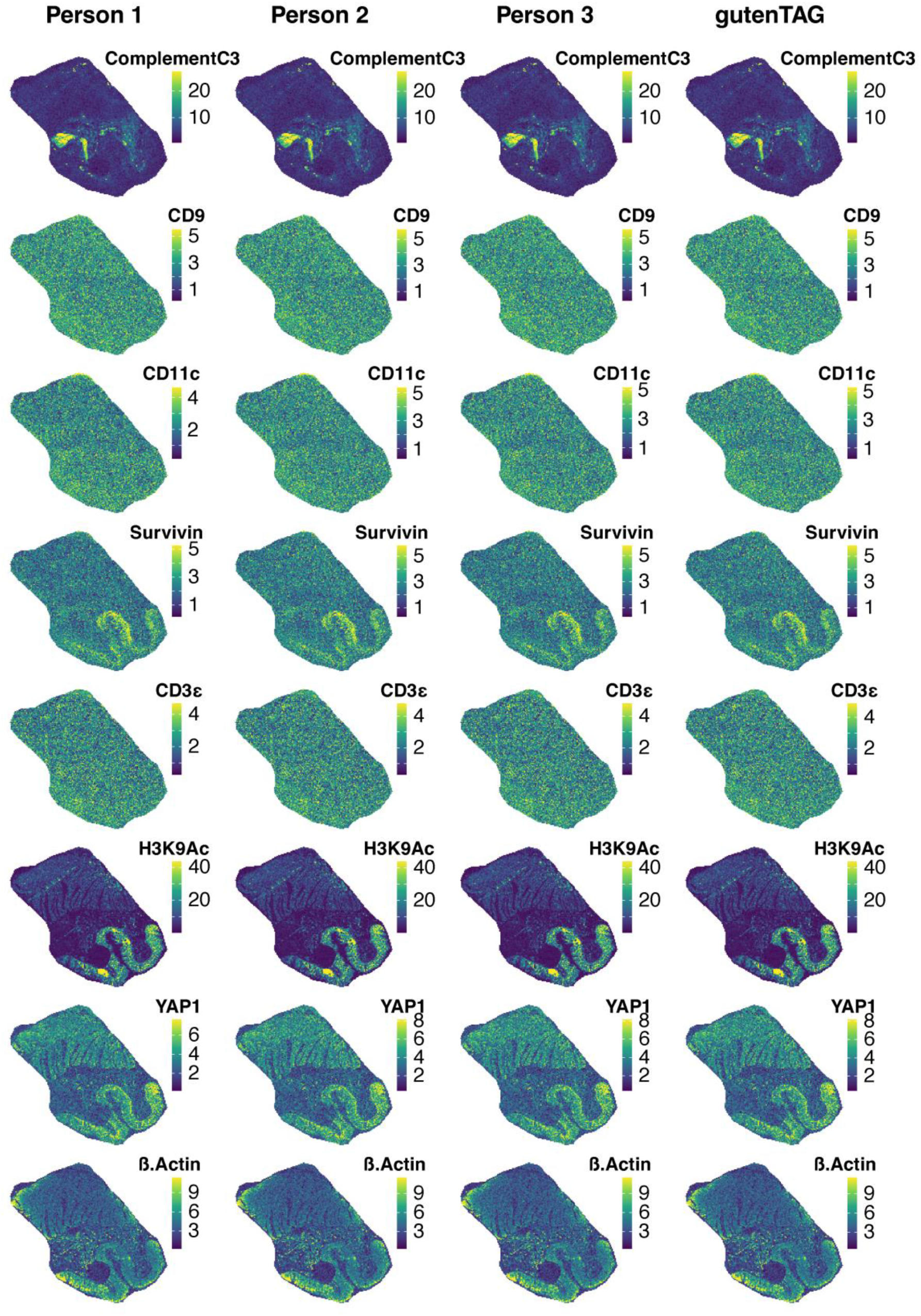
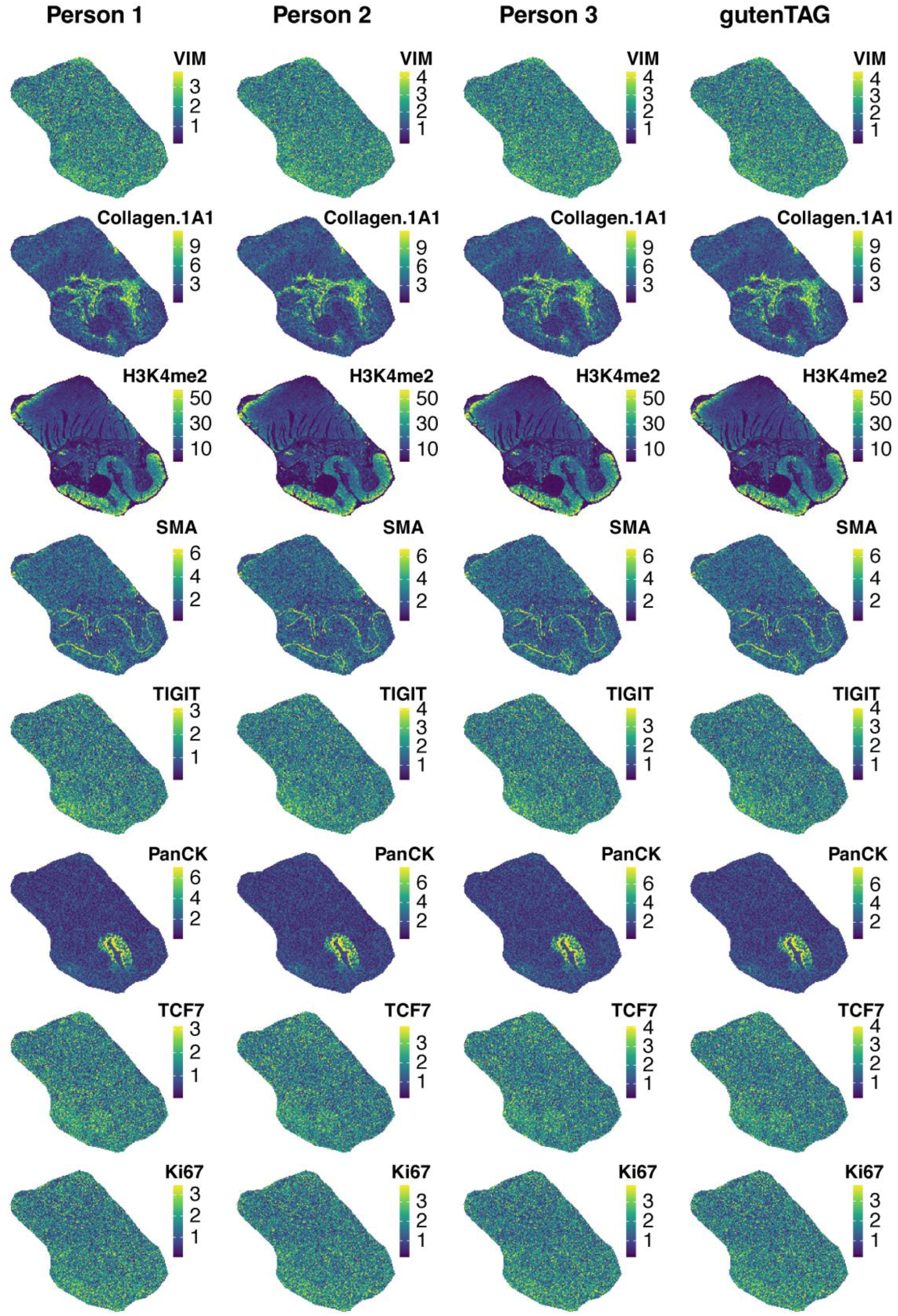
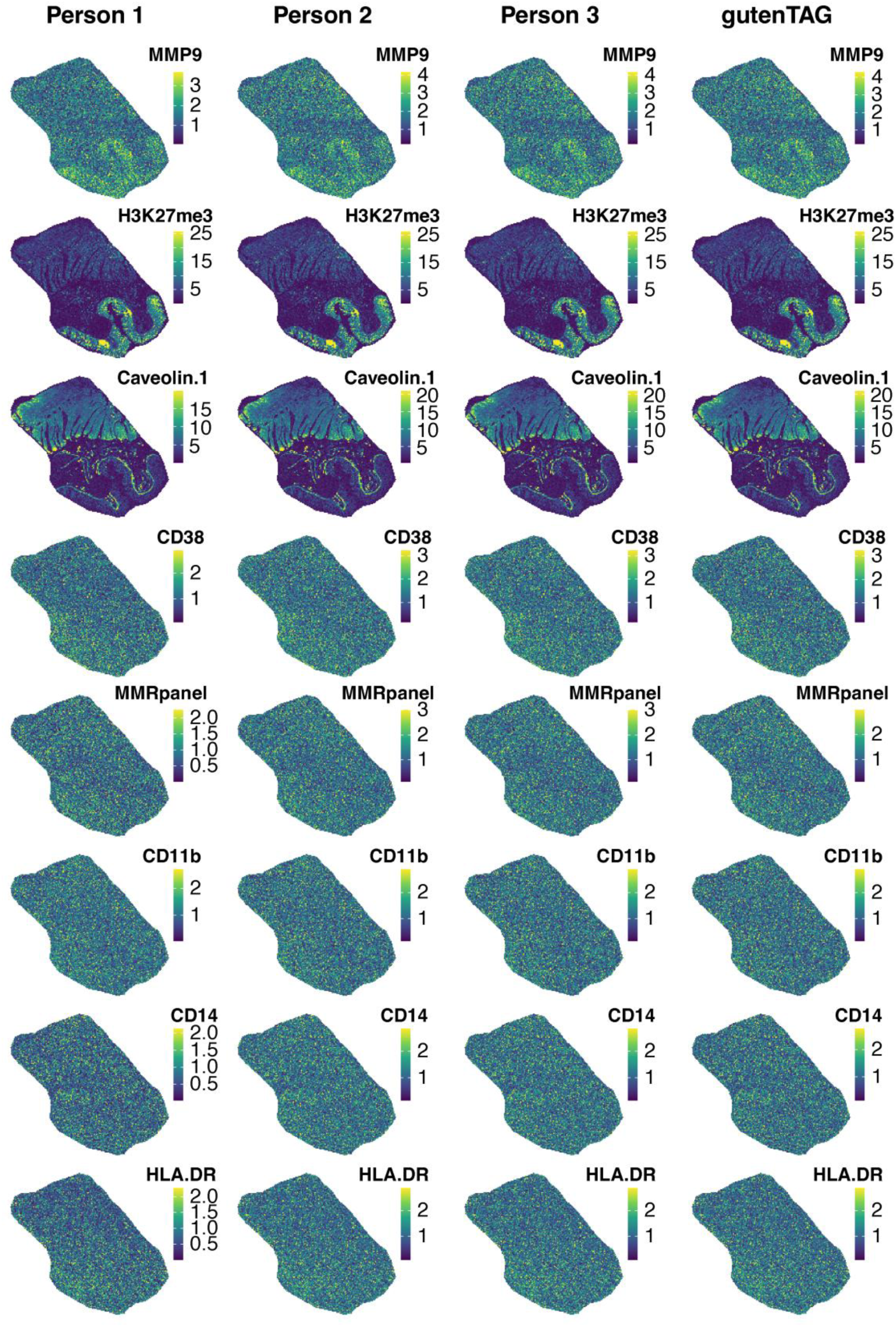
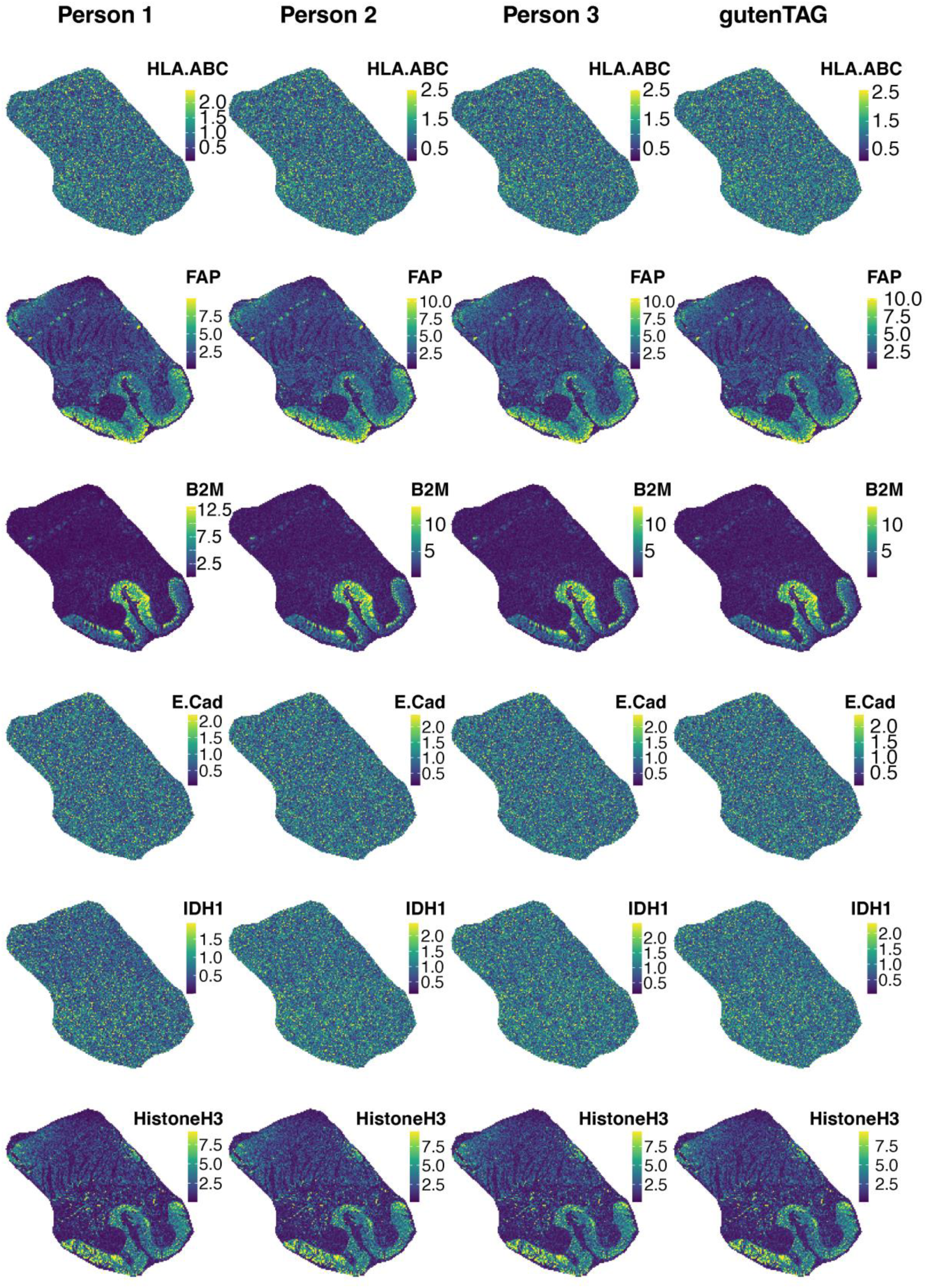
Manually picked images for all 46 PCMTs for each individual are in the three leftmost columns. In the column on the right are the gutenTAG-picked images.

**Supplementary Figure 5.**
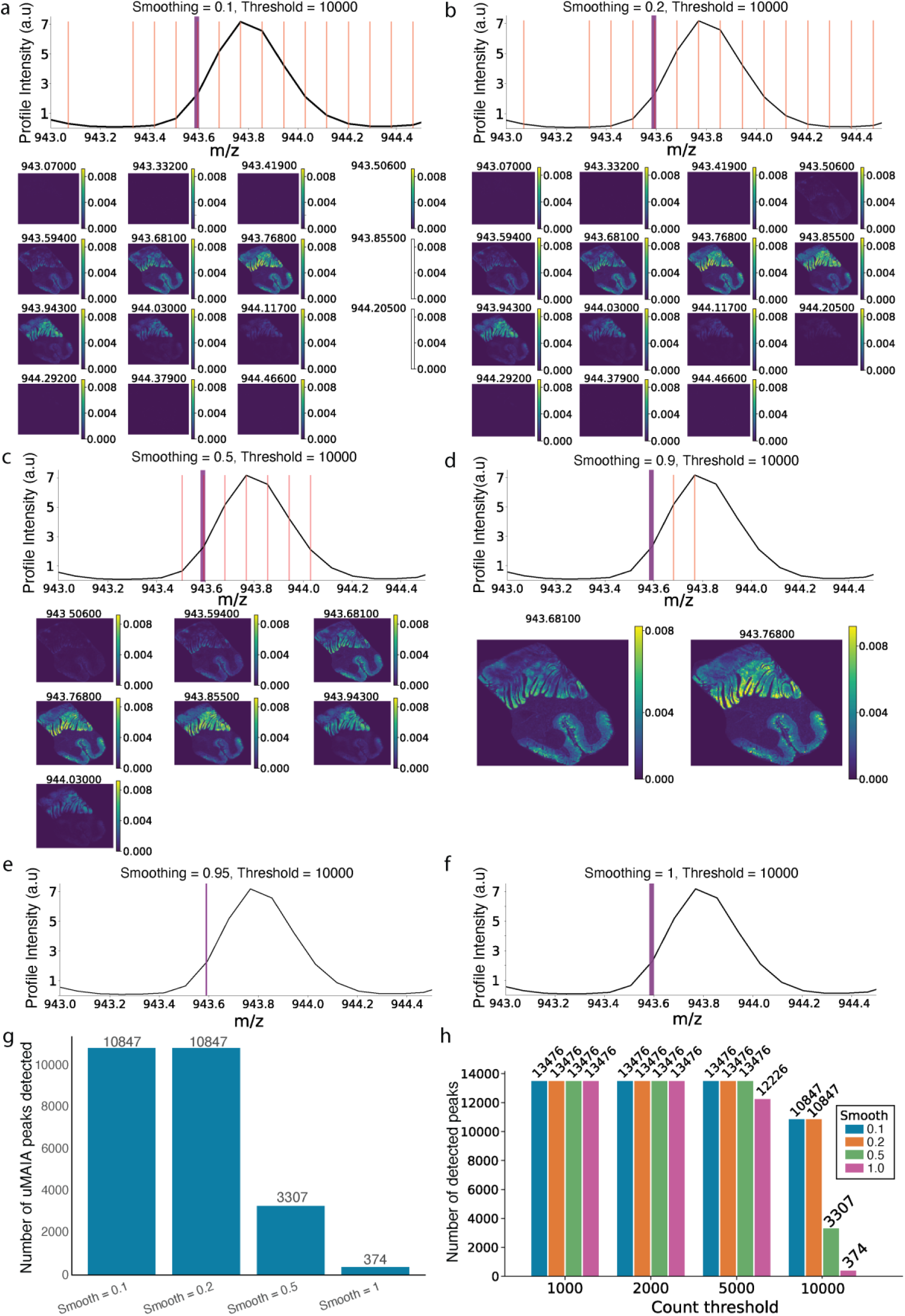
Comparison of uMAIA smoothing parameter values for peak picking in MALDI-IHC data. a-f. uMAIA peak picking using the indicated smoothing values of a) 0.1, b) 0.2, c) 0.5, d) 0.9, e) 0.95, and f) 1.0. In each case, the black line is the mean spectrum. The purple line is the expected m/z location of the CD98 mass tag. Red lines indicate the m/z values of uMAIA-picked peaks. Below are the ion images corresponding to each uMAIA peak. In each case, a count threshold of 10,000 pixels was used. No images are shown for panels e and f, as no peaks were picked. All intensity scales for images are in arbitrary units (a.u). g. The number of peaks detected at different smoothing parameter values. h. Number of peaks detected at different count threshold and smoothing values, grouped by count threshold.

**Supplementary Figure 6.**
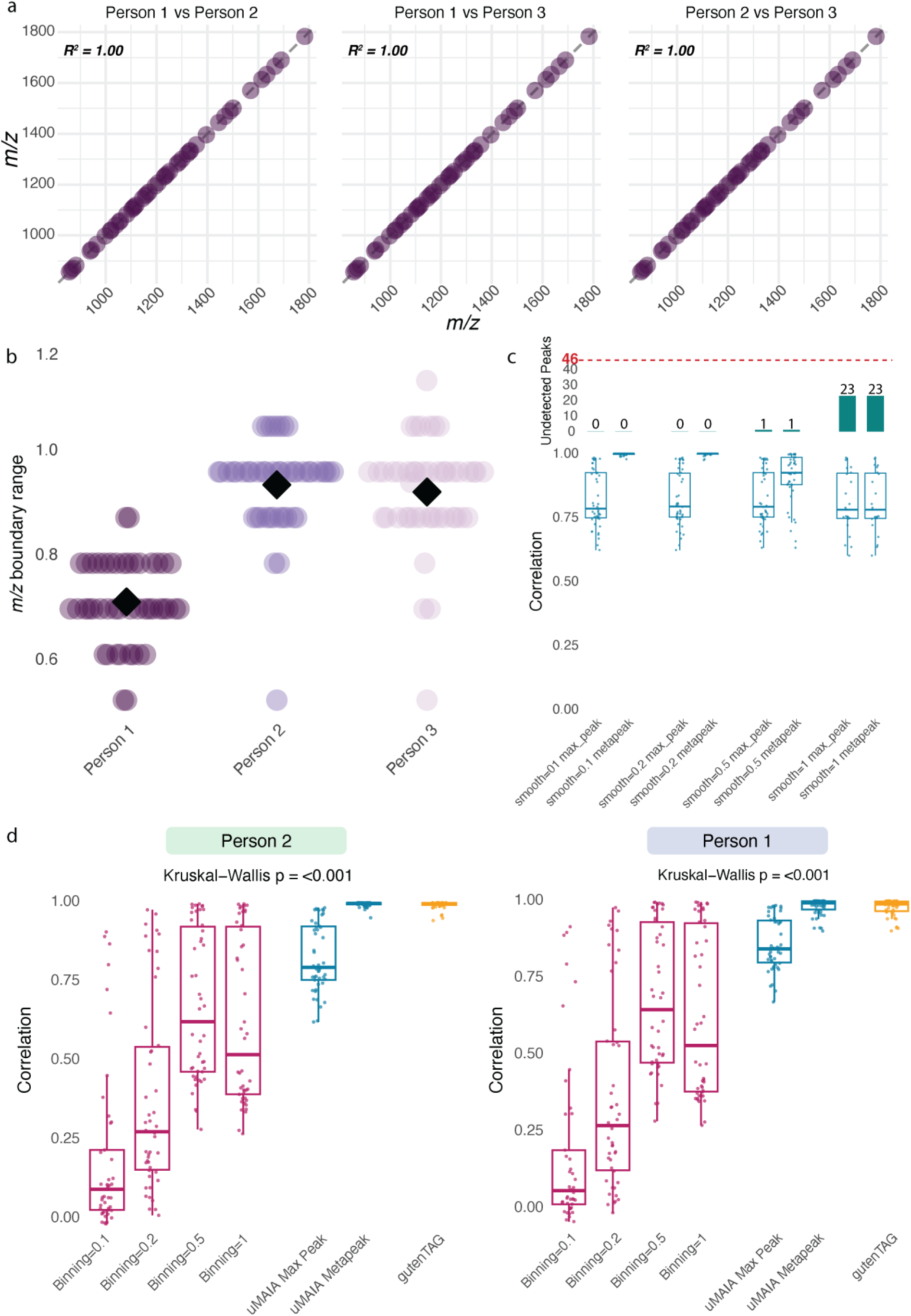
Correlation to manual images. a. Manually picked peak centers across individuals. b. Manually picked peak boundary ranges for each of the users. c. Correlations between manually picked images and images based on uMAIA under indicated conditions. The correlation to the manual image for the max intensity uMAIA peak and the uMAIA metapeak with limits = 0.5 m/z is shown for each smoothing value. Above the boxplot is the number of undetected PCMTs for each test. d. Correlations of each method to the manually picked single channel images from persons 1 and 2.

## Discussion

Use of mass spectrometry in imaging has rapidly evolved. Until recently, untargeted applications such as lipidomics and metabolomics have dominated the field; however, with the advent of probes such as PCMTs, mass spectroscopy can now be used in targeted analyses of proteins. The highly multiplexed nature of MALDI-IHC necessitates the development of computational tools designed specifically for processing and analysis of these targeted imaging experiments. We developed gutenTAG, an R-based, open-source data processing and QC tool for MALDI-IHC data. We demonstrate here that gutenTAG is a fast, high-fidelity, and scalable tool suitable for processing MALDI-IHC datasets.

gutenTAG builds on the data handling capabilities of Cardinal, a popular R-based platform for processing MALDI imaging data. In a novel step, gutenTAG employs a count-based grouping of signal from a PCMT into a single metapeak, enabling automated discovery of consensus peak locations for a PCMT in data across all pixels. gutenTAG further integrates several QC measures. gutenTAG was designed to allow the user flexibility in choice of parameters. We anticipate that MALDI-IHC will become broadly adopted and, therefore, provided users with exporter functions that will allow subsequent analysis in tools including Cardinal, SpatialExperiment, AnnData, and CytoImageList.

gutenTAG’s metapeak-generation algorithm consists of four major steps. The first is construction of a counts histogram based on compiled locations of detected peaks for all channels. Second is smoothing the counts histogram. In the third step, metapeaks are constructed via watershed segmentation. Fourth, metapeaks are assigned to the reference panel of mass tags. We have extensively characterized the effects of the parameters that control the outputs of each step. To promote transparency in peak picking, gutenTAG has interactive plotting tools that can be used to evaluate metapeak picking after each step. The optimal parameters depend on the sample and the instrument used. Therefore, data exploration and empirical testing of different parameter values is recommended. Parameter selection comes with tradeoffs, and selection of parameters that are too high or two low induces artifacts.

The window width of the noise estimator and SNR shape the counts histogram and, therefore, all downstream analyses. Values for these parameters should be based on observed noise levels in the spectra. The smoothing parameter impacts metapeak segmentation: Excessive smoothing risks under-segmentation, which results in unwanted signal in the metapeak, whereas insufficient smoothing causes over-segmentation and excessive numbers of metapeaks. The sparsity parameter controls the spacing of metapeaks during segmentation. A sparsity parameter value that is too low results in large numbers of metapeaks in close proximity, resulting in duplicate assignments that the user must then parse. In contrast, sparsity parameter values that are too high increase the number of false negatives. We found it best to err on the side of small sparsity values and manually evaluate duplicated metapeaks to find the correct match. Removal of duplicates can be automated with a gutenTAG function if the user has an expectation of the direction and extent of mass shift in their data. The metapeak limits parameter controls the lateral inclusion of signal into the metapeak. Here we suggest choosing a value that spans from the center of the metapeak to the closest local minimum on the counts histogram, as this will capture the signal from that peak in all pixels. It is also possible to include isotopic peak signal in a metapeak by using the automatic and unconstrained watershed segmentation boundaries. This can, however, introduce unwanted signal from neighboring non-isotopic peaks, especially in noisy or low-count regions of the spectrum.

To explore the parameter space, we recommend that the user employ down sampling to expedite the search process. We recognize that the need for manual parameter optimization is a limitation of the tool in its current form and aim to improve this with automated parameter estimation methods in future. We also expect that some acquisition artifacts in the data that gutenTAG was designed to correct, such as mass shifts and varying peak locations across pixels, will be mitigated as instruments improve in mass accuracy and resolution.

Dataset size is expected to increase rapidly in the coming years as MALDI-IHC experiments move to higher multiplexity, larger tissue areas, smaller pixel sizes, and larger cohorts. Therefore, speed and scalability are of crucial importance. The speed of gutenTAG is comparable to a simple approach of binning and is much faster than uMAIA, a peak picking method designed for spatial lipidomics and metabolomics. Further, gutenTAG picks peaks with higher fidelity than simple binning when assessed against the gold standard of manually picked peaks. Both uMAIA and gutenTAG are robust to mass shifts, and, interestingly, implementing a metapeak-like strategy in uMAIA improved peak picking performance, indicating that the metapeak approach is tool-agnostic.

In summary, in gutenTAG we have developed a fast, high-fidelity tool for processing and QC of MALDI-IHC datasets. We view such semi-automated and open-source computational resources as a necessary advancement to facilitate the wide adoption of MALDI-IHC. We prioritized a community-friendly design philosophy, emphasizing integration with existing community software platforms and transparent algorithmic processes, that does not sacrifice performance or scalability. Our hope is that gutenTAG contributes to the cultivation of a strong and vibrant open-source, MSI software community.

## Materials and Methods

### Data Acquisition

Following data acquisition, MALDI-IHC data were converted to the open-source .i*mzML* and .*ibd* formats. gutenTAG uses Cardinal to import .imzML data. Thcane gutenTAG pipeline can either be run locally or on a high-performance cluster such as the B-Fabric platform^21^ in the Functional Genomics Center Zürich. R version 4.4.1 was used for gutenTAG processing in B-Fabric. R version 4.3.1 was used for analysis.

### Pre-processing

Data normalization was performed using the TIC method to adjust for variations across pixels while preserving biological variability. Spectral smoothing was applied using a Gaussian filter to enhance signal quality by reducing noise, and baseline reduction was achieved through a local minimum approach, where the baseline was estimated within small m/z windows and subtracted to improve peak detection. Default parameters from the Cardinal R package were used for all preprocessing steps.

### Metapeak Generation

Metapeak construction begins by summing detected peaks across all pixels to generate a counts histogram. This is controlled by a noise estimation window parameter and the SNR parameter. Gaussian smoothing over the counts is then performed to create the landscape for subsequent metapeak segmentation. This step is controlled by a smoothing parameter, which is the standard deviation of a Gaussian filter. A watershed algorithm from the R package EBImage was then applied to the smoothed counts to segment metapeaks. Spectral signals that are segmented together are termed metapeaks. The spacing of metapeaks is affected by the smoothed counts landscape but is more directly controlled by the sparsity parameter. The sparsity parameter controls the spacing of segmentation seeds, which are local maxima of the smoothed counts histogram that act as initial marks for segmentation. The sparsity parameter therefore influences the number of metapeaks formed as well as their lateral spacing on the m/z axis.

A metapeak may result from a single peak or multiple peaks clustered together. Clustering is impacted by the value chosen for the metapeak limits parameter. The scope of a metapeak is defined by its center and its delimitations. The metapeak center is the m/z value with the highest peak counts within the limits of that metapeak; the center acts as a consensus value for the location of the majority of the peaks. Metapeak limits are the outer boundaries of a metapeak. The metapeak limits parameter can be determined automatically based on watershed segmentation or can be set manually by the user.

Finally, metapeaks are assigned to the reference panel of mass tags. Metapeak centers within a user-definable threshold of a PCMT in the panel are annotated with that tag label. Metapeaks that are outside of this association threshold are stored together with the annotated ones in a separate dataframe called AllMetapeaks. Large association thresholds (> 0.1m/z) are feasible only

### Quality Control Metrics

#### SNR calculation

The SNR of the single channel images is derived via a Gaussian mixture model to classify pixel intensities of a given channel as either signal or noise. We define SNR as the ratio between the mean intensity of these signal and noise clusters. The R package mclust was used for this calculation.

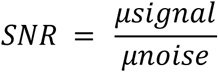

#### Geary’s C calculation

Geary’s C was computed to quantify spatial autocorrelation, calculated as:

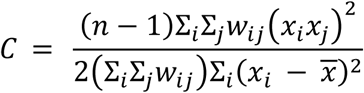

where *w_ij_* is a spatial weight matrix, *x_i_* is the intensity value in pixel i, *x_j_* is the intensity value in pixel j, *x̅* is the mean intensity value, and Σ*_i_*Σ*_j_w_ij_* is the sum of all values in the spatial weight matrix.

#### Integration of SNR and Geary’s C

SNR and Geary’s C scores for each marker are taken together to form a biaxial scatter plot that effectively describes the quality of each single channel image in terms of intensity and spatial distribution.

#### Mean-variance analysis

Mean mass tag intensities were plotted against their standard deviation and a linear regression was computed. Quantities were log scaled for variance stabilization. Standard deviation is used for plotting. Residuals were taken from the log-log mean standard deviation plot.

#### Otsu thresholding on targeted TIC

Otsu’s method is used for determining the threshold for total ion current (TIC) histogram of the targeted spectrum. For the purpose of QC, we define TIC as the sum of all assigned metapeaks, rather than the integration of all signal under the curve of a profile spectrum, as is typically meant. The TIC histogram typically shows a strong bi-modal distribution as it is the combined signal of all channels; it can often be used as a simple binary classifier for on-tissue and off-tissue pixels, as off-tissue pixels should be low across all channels.

### Parameter characterization

A parameter sweep was performed on a single colon tissue stained with a 46-plex panel for each of the parameters described (window, SNR, smoothing, sparsity, limits). Each parameter was altered individually, with default values chosen for the other parameters. Default values used were Window = 50, SNR = 2, Smoothing = 1, Sparsity = 1, Limits = 0.5. Two minor parameters, the threshold parameter (i.e., the minimum threshold of pixels to count as a segmentation border) and an m/z association parameter (i.e., the maximum m/z distance that results in assignment to a reference mass tag), were not included in this characterization. The threshold parameter was deemed to have limited influence on downstream results, and the m/z association parameter simply modulates the m/z distance allowing assignment of a metapeak to a reference mass tag and therefore behaves predictably and intuitively. The values for these parameters were: threshold = 0.01 (1% of pixels) and m/z association = 1.

The parameter values tested were as follows:

- Window: 1, 2, 5, 10, 20, 30, 40, 50 (m/z)
- SNR: 1, 2, 3, 4, 5
- Smoothing: 0.1, 0.3, 0.5, 1, 2
- Sparsity: 1, 2, 3, 4, 5, 10
- Limits: 0.1, 0.25, 0.5, 1, 2

These values were chosen to span reasonable upper and lower bounds based on experience with MALDI-IHC data. It is important to note that with a different instrument, a different range of values may be optimal.

### Method Comparison

#### Binning methods

Pre-processed .imzML and .ibd files for the 46-plex stained colon section were read into R using Cardinal’s readMSIData function. First, pre-processed profile spectra were recalibrated by using the estimateReferencePeaks() function and the recalibrate() function. Recalibration was performed using the locmax method with a tolerance of 0.1 m/z. Peak picking was then performed on the recalibrated profile data using the peakPick() function. MAD peak picking was used with SNR of 2 and window width of 50 m/z. Next, the centroided picked data were aligned using the peakAlign() function, and these data were binned with a binning resolution of 1 m/z using Cardinal’s bin() function.

#### uMAIA

Prior to running the uMAIA pipeline, data were pre-processed as above. The same data subsets (10 to 100,000 pixels) were generated to create the successive pixel size dataset. The uMAIA pipeline consisted of defining the key parameter values and running the uMAIA.pf.run() function on each dataset. The spectrum range was defined as 800-2000 m/z, and a smoothing value of 0.1 was used. Smoothing values of 0.1, 0.2, 0.5, 0.9, 0.95, and 1 were tested with a count threshold value of 10000. Count thresholds of 1000, 2000, 5000, and 10000 were tested.

### Statistics

To assess the correlations between automatic and the ground truth manual peak picking images, the Pearson correlation coefficient between the method’s intensity vector and the corresponding manually picked reference vector was computed (cor(), R stats package, with use = pairwise.complete.obs to handle missing pixels). This yielded one correlation coefficient per peak per method, quantifying how faithfully each method reproduced the spatial intensity distribution of the manual reference. Four fixed-width binning approaches (bin widths 01, 0.2, 0.5, and 0.1 Da) were used for Cardinal’s binning. Two uMAIA configurations were used: the maximum intensity peak within a distance of 0.5 m/z on either side of the reference mass tag (Max Peak Only), and a metapeak strategy that integrated signal from all uMAIA peaks within 0.5 m/z on either side of the peak with the maximum intensity within a proximity of 0.5 m/z of the reference peak. Differences in the resulting correlation distributions across methods were assessed using a Kruskal–Wallis rank-sum test. The *p* values comparing peak centers of the peak picking methods to the manually-picked peak centers were calculated using a Benjamini-Hochberg-corrected *t*-test.

## Data and Code Availability

The gutenTAG repository is available at https://github.com/BodenmillerGroup/gutenTAG. The code for generating the figures for the paper will be available upon publication.

## Acknowledgements

We acknowledge the help of Lasse Meyer, Quentin Hellier, and Lydia Schulla in performing the manual peak picking. We appreciate the support of the members of the proteomic facility at FGCZ.

